# Consolidated Bioprocessing of Lignocellulosic Biomass Poplar to Produce Short-Chain Esters by *Clostridium thermocellum*

**DOI:** 10.1101/2023.03.29.534841

**Authors:** Hyeongmin Seo, Priyanka Singh, Charles E. Wyman, Charles M. Cai, Cong T. Trinh

## Abstract

Consolidated bioprocessing (CBP) of lignocellulosic biomass using cellulolytic microorganisms presents a promising sustainable and economical biomanufacturing platform where enzyme production, biomass hydrolysis, and fermentation to produce biofuels, biochemicals, and biomaterials occur in a single step. However, understanding and redirecting metabolism of microorganisms to be compatible with CBP to produce non-native metabolites are limited. In this study, we metabolically engineered a cellulolytic thermophile *Clostridium thermocellum* and demonstrated its compatibility with CBP integrated with a mild Co-solvent Enhanced Lignocellulosic Fractionation (CELF) pretreatment for conversion of hardwood poplar into short-chain esters (i.e., ethyl acetate, ethyl isobutyrate, isobutyl acetate, isobutyl isobutyrate) with broad use as solvents, flavors, fragrances, and biofuels. A recombinant *C. thermocellum* engineered with deletion of carbohydrate esterases and stable overexpression of a thermostable alcohol acetyltransferase improved the target esters production without compromised deacetylation activities. We discovered these esterases exhibited promiscuous thioesterase activities and their deletion improved ester production by increasing isobutanol flux and rerouting the native electron and carbon fermentative metabolism besides their known major function of ester degradation. The total ester production could be further enhanced up to 80-fold and the composition of short-chain esters could be modified by deleting lactate biosynthesis and/or CELF-pretreated poplar under different pretreatment conditions.

## Introduction

Microbial biosynthesis enables an economical conversion of lignocellulose (e.g., woody biomass) into renewable biofuels, biochemicals, and biomaterials with high selectivities under mild conditions^1, 2^. Due to complex multi-scale plant cell well structure, this bioconversion of recalcitrant lignocellulose, however, often needs energy-extensive pretreatment processes and the addition of costly enzymes to aid in hydrolyzing polysaccharides into fermentable monosaccharides^3^. Consolidated bioprocessing (CBP) eliminates enzyme addition prior to the fermentation by employing cellulolytic microorganisms, capable of producing cellulolytic enzymes, solubilizing and fermenting polysaccharides to produce a target biochemical^4, 5^. To maximize economic advantages of CBP, an integrative design of the mild ‘upstream’ feedstock process engineering and the ‘downstream’ strain development and fermentation is important^6^.

Co-solvent Enhanced Lignocellulosic Fractionation (CELF) pretreatment is a mild biomass preprocessing step that can be beneficial for CBP^7^. CELF is compared favorably with other leading pretreatment technologies, particularly when producing pretreated biomass solids for bioconversion^8^. CELF pretreatment uniquely employs an aqueous mixture containing water, tetrahydrofuran (THF), and dilute acid to enhance the solubilization of hemicellulose and lignin from cellulose in biomass and to promote greater micro- and macro-accessibility of the pretreated solids to enzymatic accessibility^9–11^. In studies pairing CELF with enzymatic hydrolysis or Simultaneous Saccharification and Fermentation (SSF), CELF-pretreated biomass solids were found to be far more conducive to enzymatic breakdown, requiring 90% less enzyme loadings than dilute-acid pretreated biomass, achieving nearly complete solubilization of solids when incubated with a cellulolytic microorganism^12^. Integration of CELF pretreatment in CBP can be potentially transformative for bioconversion of lignocellulosic biomass to produce biofuels and chemicals but is underexplored. Optimization of CELF pretreatment reaction conditions can further lead to enhanced physiochemical changes to the biomass for enhanced deconstruction by CBP^13^. In addition, upgrading the CELF reactor to a larger scale unit that more closely resembles a commercially relevant process could further improve digestibility of the pretreated biomass. Therefore, co-optimization of CELF with CBP utilizing microbial biocatalysts for tailored biochemical and biofuel production has enormous potential.

Conventional CBP of biomass has been studied for production of native products such as ethanol that is inherent to fermentative metabolism of cellulolytic microorganisms. For instance, a cellulolytic thermophile *Clostridium thermocellum* (also known as *Acetivibrio thermocellus*) is a representative CBP microbe exhibiting exceptional capability of lignocellulose solubilization and fermentation for ethanol production^5, 14^. The endogenous metabolism of *C. thermocellum* can be rewired to produce isobutanol^15^ and butanol^16^, diversifying product molecules from lignocellulose fermentation, an important innovation required for advanced CBP technology^6^. Promising pivot products of CBP are short-chain esters^17^, which are naturally found in ripening fruits and flowers. Alcohol acyltransferase (AAT), condensing an alcohol and an acyl-CoA to form an ester, is a major ester biosynthesis enzyme commonly found from the natural ester-producing plants^18^ or yeasts^19^. The AAT-dependent pathways can be harnessed to synthesize a large space of designer bioesters including acetate esters^20–28^, propionate esters^22^, lactate esters^20, 29^, butyrate esters^21, 28, 30^, pentanoate esters^22^, and hexanoate esters^22^. Because eukaryotic AATs known to date are not thermostable and poorly expressed in a foreign host^31–33^, they are incompatible with elevated growth temperatures of *C. thermocellum* (>50℃)^34^. Our recent studies demonstrated that engineering promiscuity of thermostable chloramphenicol acetyltransferases (CATs) to function as AATs enabled short-chain ester production including isobutyl acetate from *C. thermocellum*^29, 34^. A single mutation in CATec3 Y20F not only improved its thermostability by increasing the enzyme melting temperature by ∼7°C, but also enabled it to synthesize a large space of short-chain esters, at least 168 ester molecules with various combinations of 8 acyl-CoAs and 21 alcohols^29^. Successful application of the robust AATs in the CBP microorganisms can be potentially transformative by expanding their metabolic capabilities to produce designer bioesters as industrial platform chemicals with versatile use for food additives^35^, green solvents^36^, and potential drop-in biofuels^37^ (Fig. 1). However, *C. thermocellum* compatibility with CBP to produce non-native compounds such as esters is poorly understood.

**Figure 1.**
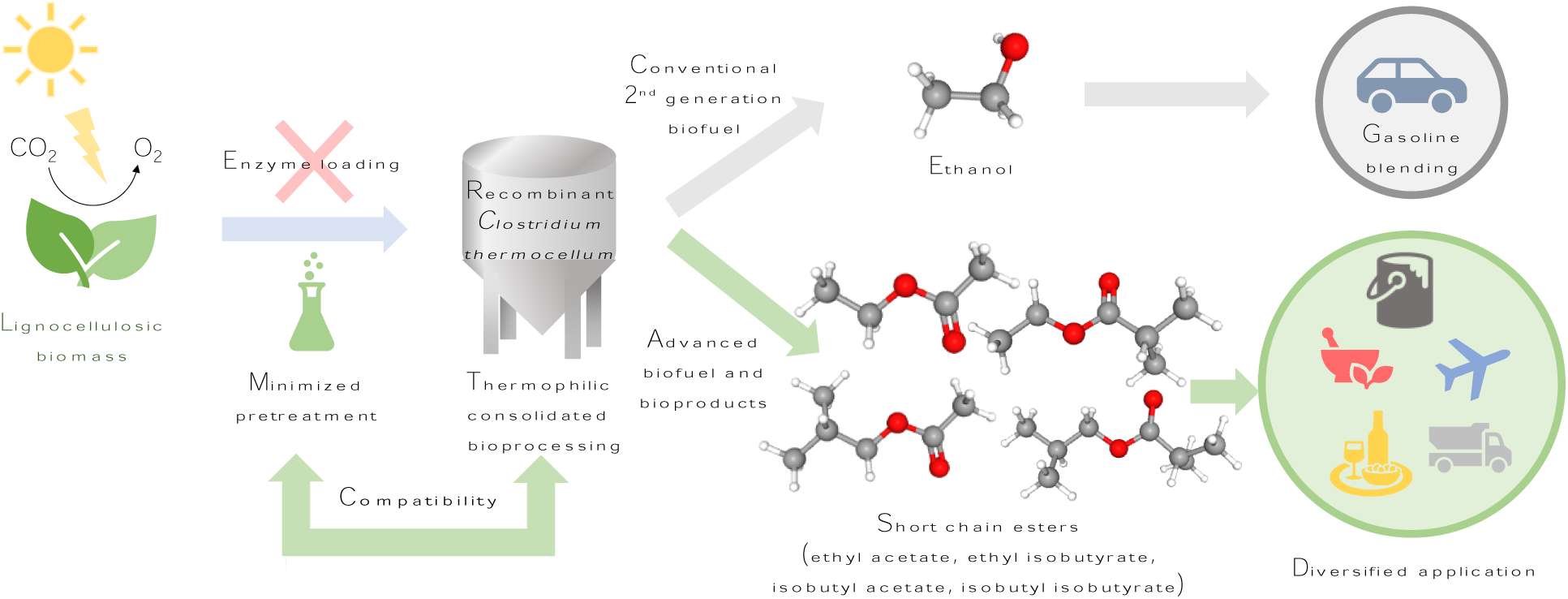
A scheme of consolidated bioprocessing of lignocellulosic biomass for biosynthesis of non-native short-chain (C4-C8) esters using a cellulolytic thermophile *C. thermocellum*.

In this study, we investigated compatibility of *C. thermocellum* with CBP that was integrated with CELF pretreatment for direct conversion of CELF-pretreated woody poplar to short-chain (C4-C8) esters, including ethyl acetate, ethyl isobutyrate, isobutyl acetate, and isobutyl isobutyrate. These esters are derived from the endogenous acetyl-CoA and isobutyl-CoA overflow fermentative metabolism of *C. thermocellum.* We first elucidated how the underground metabolism of a cellulolytic thermophile *C. thermocellum* could be modulated to activate the short-chain ester biosynthesis. We next examined the effect of manipulating carbon and electron competing pathways (i.e., deletion of carbohydrate esterases and lactate biosynthesis and overexpression of thermostable CATs) on the solubilization of CELF-pretreated poplar, redox metabolism, and production of short-chain esters. Finally, we interrogated the impact changes to CELF pretreatment reaction conditions with two different reactor types on the redox metabolism of *C. thermocellum* and its conversion of CELF-pretreated poplar to the target esters.

## Results and Discussion

### Activating the endogenous metabolism of *C. thermocellum* for short-chain ester biosynthesis

#### Pathway design

Even though wildtype *C. thermocellum* does not natively produce C4-C8 short-chain esters (e.g., ethyl acetate, ethyl isobutyrate, isobutyl acetate, and isobutyl isobutyrate), it has endogenous metabolism to produce the precursor metabolites acetyl-CoA, isobutyryl-CoA, ethanol, and isobutanol for biosynthesis of these target esters (Fig. 2A). Due to optimal growth of *C. thermocellum* at elevated temperatures (>50℃) and its unique capability to make cellulolytic enzymes (e.g., esterases) to degrade lignocellulosic biomass, a combined heterologous expression of a thermostable AAT and deletion of the select endogenous ester-degrading carbohydrate esterases (CEs) are required to activate the short-chain ester biosynthesis pathways in *C. thermocellum*^29, 38^. Since the biosynthesis of the target alcohols and acyl-CoAs is important for the anabolic and redox metabolisms of *C. thermocellum*, which is responsive to both genetic and environmental perturbations^39–43^, we first investigated how these two genetic manipulations activated the biosynthesis of short-chain esters and affected cell growth, biomass degradation, and redox metabolism using the model cellulose substrate Avicel, which is a commercial, pure crystalline cellulose derived from wood pulp.

**Figure 2.**
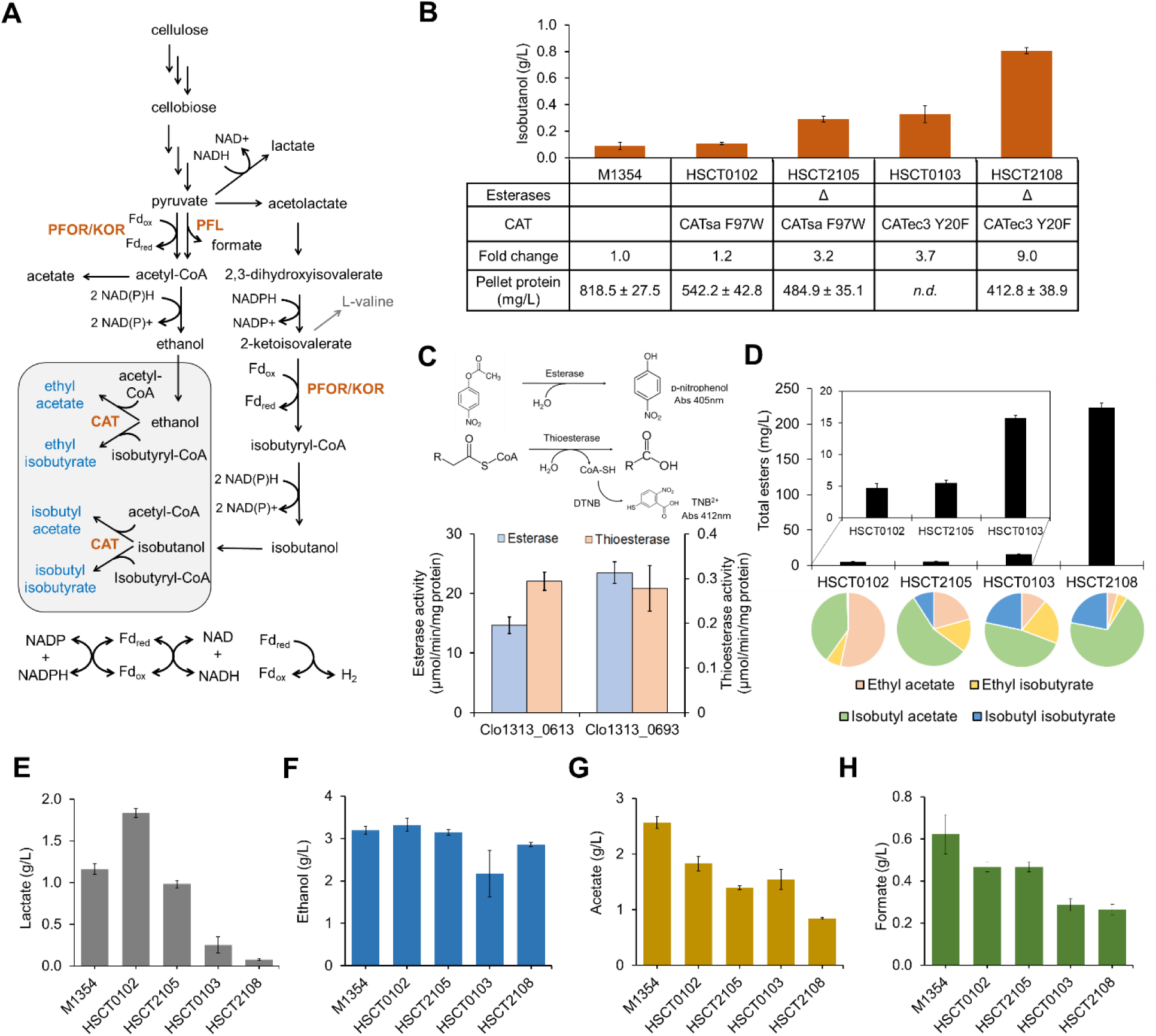
Harnessing the endogenous acetyl-CoA and isobutyryl-CoA metabolism of *C. thermocellum* for consolidated bioprocessing of lignocellulosic biomass to produce non-native short-chain esters. **(A)** Metabolic pathways for short-chain ester biosynthesis. **(B)** Comparison of isobutanol production by engineered *C. thermocellum* strains. **(C)** Promiscuous thioesterase activities of the two carbohydrate esterases Clo1313_0613 and Clo1313_0693. **(D-H)** End-point titers of **(D)** short-chain esters, **(E)** lactate, **(F)** ethanol, **(G)** acetate, and **(H)** formate by the engineered *C. thermocellum* strains after 72 h Avicel fermentation. Each data point represents a mean ± 1 standard deviation from at least three biological replicates.

#### Deletion of carbohydrate esterases increased isobutyryl-CoA flux, enhancing both isobutanol and ester production

We started by analyzing the metabolite profiles of the engineered *C. thermocellum* strains overexpressing thermostable CATs either with or without deletion of the two CE-encoding genes (i.e., *clo1313_0613* and *clo1313_0693*) after 72-hour Avicel fermentation. Unexpectedly, we found that deletion of the two CEs improved isobutanol production (Fig. 2B). The esterase-deficient strain HSCT2105 produced 0.29 g/L isobutanol, about 3.0-fold higher than the parent strain HSCT0102 (0.11 g/L) without two CE deletions^38^. Likewise, the esterase-deficient strain HSCT2108 produced 0.81 g/L of isobutanol, about 2.5-fold higher than the parent strain HSCT0103 (0.33 g/L). There were no significant differences in cell pellet proteins (indirect measurement of cell mass) between the engineered strains. These results suggested that deletion of the two CEs enhanced metabolic fluxes towards the isobutanol biosynthesis.

While the effects of deleting these two CEs on lignocellulose degradation and short-chain ester hydrolysis were previously reported^38, 44^, it was still unknown of how their deletion improved both the target alcohol and ester production besides their main function. Since many esterases also function as phospholipases, thioesterases, and/or proteases^45, 46^, we hypothesized that the two CEs might have exhibited promiscuous activities affecting the endogenous isobutyryl-CoA metabolism. Because the isobutanol production pathway of *C. thermocellum* is derived from isobutyryl-CoA (Fig. 2A), we next examined whether the two CEs have thioesterase activities that hydrolyze isobutyryl-CoA. The *in vitro* enzyme characterization showed that both CEs indeed had minor thioesterase activities against isobutyryl-CoA (Fig. 2C). The thioesterase activities of Clo1313_0613 and Clo1313_0693 were 50- and 84-fold lower than the esterase activities of Clo1313_0613 and Clo1313_0693, respectively. Because the intracellular level of isobutyryl-CoA was low in a micromolar concentration range, deletion of the two CEs might have significantly increased isobutyryl-CoA availability used for the isobutanol and short-chain ester production.

#### Thermostable AATs improved biosynthesis of short-chain esters with higher isobutanol production and subsequent alcohol conversion

Since the AAT-dependent ester biosynthesis utilizes acyl-CoAs and alcohols as substrates, the more robust and efficient AATs are expected to enhance the turnover rates of these substrates in *C. thermocellum*^29^. To investigate how the expression of AATs could affect the availability of acyl-CoAs and alcohols and alter the endogenous metabolism of *C. thermocellum*, we compared the isobutanol production by the strains expressing CATs that exhibit different thermostability and catalytic efficiency towards isobutanol^29, 47^. When CATsa F97W was overexpressed in HSCT0102, the isobutanol production reached up to 0.11 g/L, which was ∼1.2-fold higher than the control strain M1354 (0.08 g/L) (Fig. 2B). When a more thermostable and efficient AAT (CATec3 Y20F) was overexpressed in HSCT0103, the isobutanol production was 3.7-fold higher than M1354. Finally, the CE-deleted strain HSCT2018 overexpressing CATec3 Y20F produced 9.0-fold higher isobutanol than M1354, suggesting that there was a synergistic effect between the CE deletion and AAT overexpression on the alcohol biosynthesis. The enhanced isobutanol production improved short-chain ester titers and changed the ester composition (Fig. 2D). Particularly, HSCT2108 produced 0.22 g/L of short-chain esters, which was about 47-fold higher titers than HSCT0102 (0.005 g/L), the ester producing *C. thermocellum* strain first ever reported^47^. Interestingly, 91% (mol/mol) of these esters produced by HSCT2108 were isobutyl esters (i.e., isobutyl acetate and isobutyl isobutyrate), while the percentage of these isobutyl esters synthesized by HSCT0102 was only 40%. Although CATec3 Y20F is known to prefer acetyl-CoA to isobutyryl-CoA as an acyl-CoA substrate^29^, the higher availability of isobutyryl-CoA in the engineered strains significantly increased the composition of ethyl and isobutyl isobutyrates (Fig. 2D). For example, HSCT2108 produced isobutyl esters up to 26% (mol/mol) of the total short-chain esters, while HSCT0102 produced isobutyryl esters only at a 6% (mol/mol) fraction. HSCT0103 achieved the highest composition (42%) of isobutyrate esters, indicating that the intracellular acetyl-CoA and isobutyryl-CoA pools were significantly modulated by both the CE deletion and CATec3 Y20F overexpression. The results also supported our conclusion that the CE deletion reduced thioesterase activity, resulting in an increase of isobutyryl-CoA availability that boosted the biosynthesis of isobutanol and isobutyl-CoA derived esters (Fig. 2C).

#### Biosynthesis of short-chain esters significantly affected the fermentative metabolism of C. thermocellum

We next investigated how the CE deletion and AAT overexpression affected the fermentative metabolism of *C. thermocellum* (Figs. 2E, F, G, H). We found that the lactate biosynthesis pathway of the engineered strains was significantly modulated (Fig. 2E). After the 72-hour Avicel fermentation, the best isobutanol and ester producer HSCT2108 produced 14.4-fold less lactate than the control strain M1354. The decrease in lactate production correlated with the increase in isobutanol production likely because production of these two fermentative products directly competed for both the precursor metabolite pyruvate and electrons of NAD(P)H (Fig. 2B). Deletion of CEs and overexpression of strong CATec3 Y20F synergistically contributed to significant reduction of lactate biosynthesis. Likewise, productions of acetate and formate in HSCT2108 were significantly reduced by 3.0-fold and 2.4-fold, respectively, as compared to M1354 (Fig. 2G, 2H). Since acetate biosynthesis is a major pathway to generate ATP that is used for cell growth under anaerobic conditions, we found that the lower acetate production correlated to the lower maximum cell yield of HSCT2108 than M1354 (Fig. 2B). The reduction in cell mass yield and acetate production could also be attributed to the competitive use of acetyl-CoA by HSCT2108 for ester biosynthesis. Unlike the other fermentative products, we found that the ethanol production remained similar in a range of 2.8 and 3.3 g/L regardless of the CE deletion and CAT overexpression (Fig. 2F).

Overall, the CE deletion and CATec3 Y20F overexpression activated the endogenous isobutyryl-CoA metabolism required for the biosynthesis of short-chain esters. These genetic changes affected the fermentative metabolism of *C. thermocellum* by enhancing electron availability for ester biosynthesis while reducing lactate biosynthesis, cell growth, and acetyl-CoA availability for ATP generation via acetate biosynthesis.

#### The ester-producing *C. thermocellum* HSCT2108 was compatible with the consolidated bioprocessing of CELF-pretreated poplar for biosynthesis of short-chain esters

In this study, CELF was configured to involve mild (160℃) pretreatment of poplar wood chips, employing aqueous tetrahydrofuran (THF) with dilute sulfuric acid to solubilize lignin and hemicellulose in order to recover glucan-rich solids^10, 48, 49^. Due to the efficient lignin removal and enhanced recovery of glucan in solids, CELF-pretreated poplar required relatively low enzyme loading for saccharification^50^, which can be beneficial for the CBP ester biosynthesis. Here, we investigated whether the ester-producing *C. thermocellum* HSCT2108 was compatible with CBP for converting CELF-pretreated poplar to short-chain esters (Fig. 3A).

**Figure 3.**
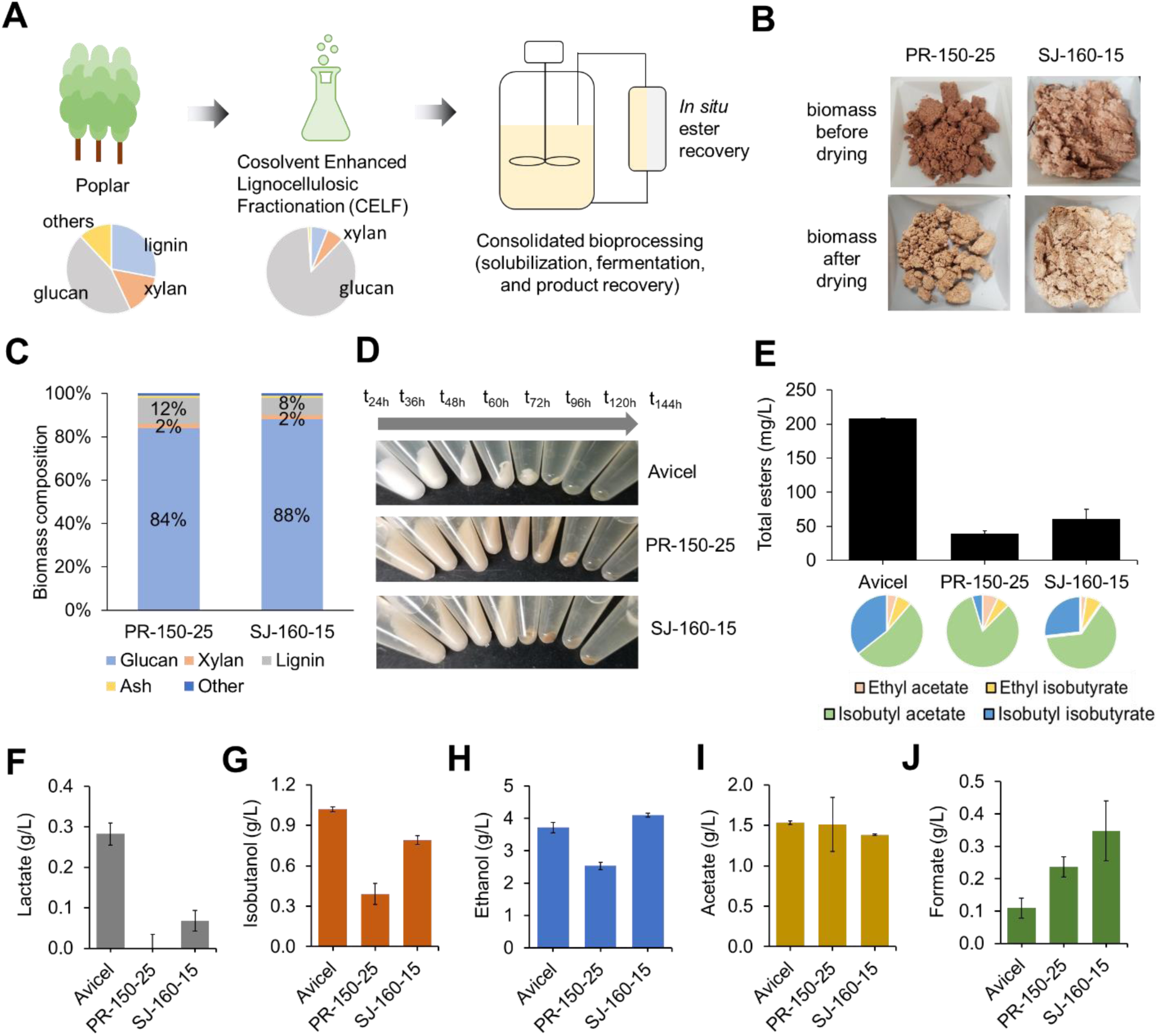
Consolidated bioprocessing of CELF-pretreated woody poplar for short-chain ester production by *C. thermocellum* HSCT2108. **(A)** A process scheme of an integrated CBP for ester production. **(B)** Images of the CELF-pretreated poplar samples processed from two different reactor runs. The PR-150-20 corresponds to the CELF process performed in a Parr reactor at 150℃ for 20 minutes while the SJ-160-15 is referred to the CELF process conducted in a Steam jacketed reactor at 160℃ for 15 minutes. **(C)** Biomass composition of the CELF-pretreated poplar. **(D)** Images of insoluble fractions collected after 144-hour fermentation using three types of lignocellulosic biomass (Avicel, CELF-pretreated poplars processed by PR-150-25 and SJ-160-15) by *C. thermocellum* HSCT2108. **(E-J)** End-point titers of **(E)** short-chain esters, **(F)** lactate, **(G)** isobutanol, **(H)** ethanol, **(I)** acetate and **(J)** formate from the three different lignocellulosic biomass fermentation by *C. thermocellum* HSCT2108. Each data point represents a mean ± 1 standard deviation from at least three biological replicates.

#### HSCT2108 solubilized woody poplar pretreated under different CELF reactor configurations

To better understand the compatibility of *C. thermocellum* with CBP coupled with CELF pretreatment for biochemical production, we examined whether HSCT2108 could grow and utilize CELF-pretreated woody poplar. We performed CELF pretreatment of poplar in a smaller scale continuously stirred 1 L Parr reactor (PR-150-25, 150℃ for 25 mins) and in a larger unstirred 3.8 L steam jacketed reactor (SJ-160-15, 160℃ for 15 mins) (Fig. 3B). The smaller scale pretreatment configuration was selected to serve as a reference point for benchmark validation comparable to previously published results from CELF at a lab scale. The larger scale configuration represents a commercially relevant setting and operating procedure that produces pretreated material that would be representative of scaled-up processing up to 300 tons per day. The CELF-pretreated poplar from the two configurations exhibited slightly different biomass compositions (Fig. 3C). The SJ-160-15 pretreatment achieved the same amount of xylan (2% w/w) but lower K-lignin contents (8% w/w) than the PR-150-25 pretreatment (12% K-lignin). The strain characterization showed that HSCT2108 could grow on woody poplar pretreated by both CELF pretreatment configurations. The CELF-pretreated poplar with a loading of ∼19 g/L glucan were completely solubilized after the 144-hour fermentation (Fig. 3D).

### Different CELF pretreatment configurations affected isobutanol and short-chain ester production by *C. thermocellum* HSCT2108

We performed the Avicel, PR-150-25, and SJ-160-15 fermentations to compare the compatibility of *C. thermocellum* HSCT2108 with CBP for short-chain ester biosynthesis. Strain characterization showed that the production of short-chain esters from the PR-150-25 (0.04 g/L) and SJ-160-15 (0.06 g/L) fermentations were 5.5- and 3.5-fold lower than the Avicel fermentation (0.21 g/L), respectively. Interestingly, the ester composition was similar between the SJ-160-15 and Avicel fermentations (Fig. 3E), suggesting that the SJ-160-15 substrate might have inhibited the metabolism of HSCT2108 less severe than the PR-150-25 substrate. The SJ-160-15 fermentation reduced lactate and isobutanol to the concentrations of 0.07 g/L and 0.79 g/L, respectively (Figs. 3F, 3G). However, the PR-150-25 fermentation produced lactate and isobutanol even at lower levels. Likewise, ethanol, a major electron sink, was produced 33% less in the PR-150-25 fermentation than in the SJ-160-15 fermentation, while both the SJ-160-15 and Avicel fermentations produced ethanol at a similar level (Fig. 3H). Acetate production did not exhibit any significant differences among the strains (Fig. 3I); however, formate production was increased by 2.1-fold and 3.1-fold for the PR-150-25 and SJ-160-15 fermentations, respectively, as compared to the Avicel fermentation (Fig. 3J).

In summary, *C. thermocellum* HSCT2108 solubilized and fermented CELF-pretreated poplar for both cell growth and short-chain ester biosynthesis, which made it compatible with CBP coupled with CELF pretreatment. However, as compared to the model cellulose substrate Avicel, both PR-150-25 and SJ-160-15 substrates were more complex due to the presence of small fractions of xylan and lignin that might have negatively interfered with *C. thermocellum* metabolism, as observed by a decrease in cell growth and biosynthesis of lactate, isobutanol and short-chain esters.

### Enhancing short-chain ester production by modulating electron and precursor availability

Since the lactate biosynthesis competes with the short-chain ester biosynthesis for pyruvate and NAD(P)H (Fig. 4A), we disrupted the lactate pathway by deleting *ldhA* (*clo1313_1160*) from the esterase*-*deficient strain HSCT2108 (Fig. 4B). Concurrently, we aimed to enhance stable CATec3 Y20F overexpression to further improve the short-chain ester production. Previous studies reported that a plasmid-based gene expression in *C. thermocellum* using an endogenous promoter could cause undesired integration of the plasmid into the genome via homologous recombination^51, 52^. The crossover event particularly occurred at the 200-400 base pair length promoter region^53^. In agreement with the previous studies, we observed that the plasmid-based overexpression of CATec3 Y20F under the 310 bp length of *gapDH* promoter (pHS0024) triggered such plasmid integration at the genomic *gapDH* promoter region after 60-hour fermentation (Fig. S3). Since the plasmid integration diminished gene copy numbers, compromising stable expression of the protein^52^, we reasoned that a more stable CATec3 Y20F overexpression would improve short-chain ester production. To reduce the frequency of homologous recombination that depends on the length of homology nucleotide sequences^54^, we truncated the *gapDH* promoter by eliminating 130 bp of the initial sequence and expressed CATec3 Y20F under the control of 180 bp length version of the *gapDH* promoter (pHS0070). The truncated *gapDH* promoter included a putative SigA/RpoD motif to ensure integrity of transcription initiation^51^. We generated HSCT3111 derived from HSCT2108 with both *ldhA* deletion and CATec3 Y20F overexpression from the pHS0070 plasmid.

**Figure 4.**
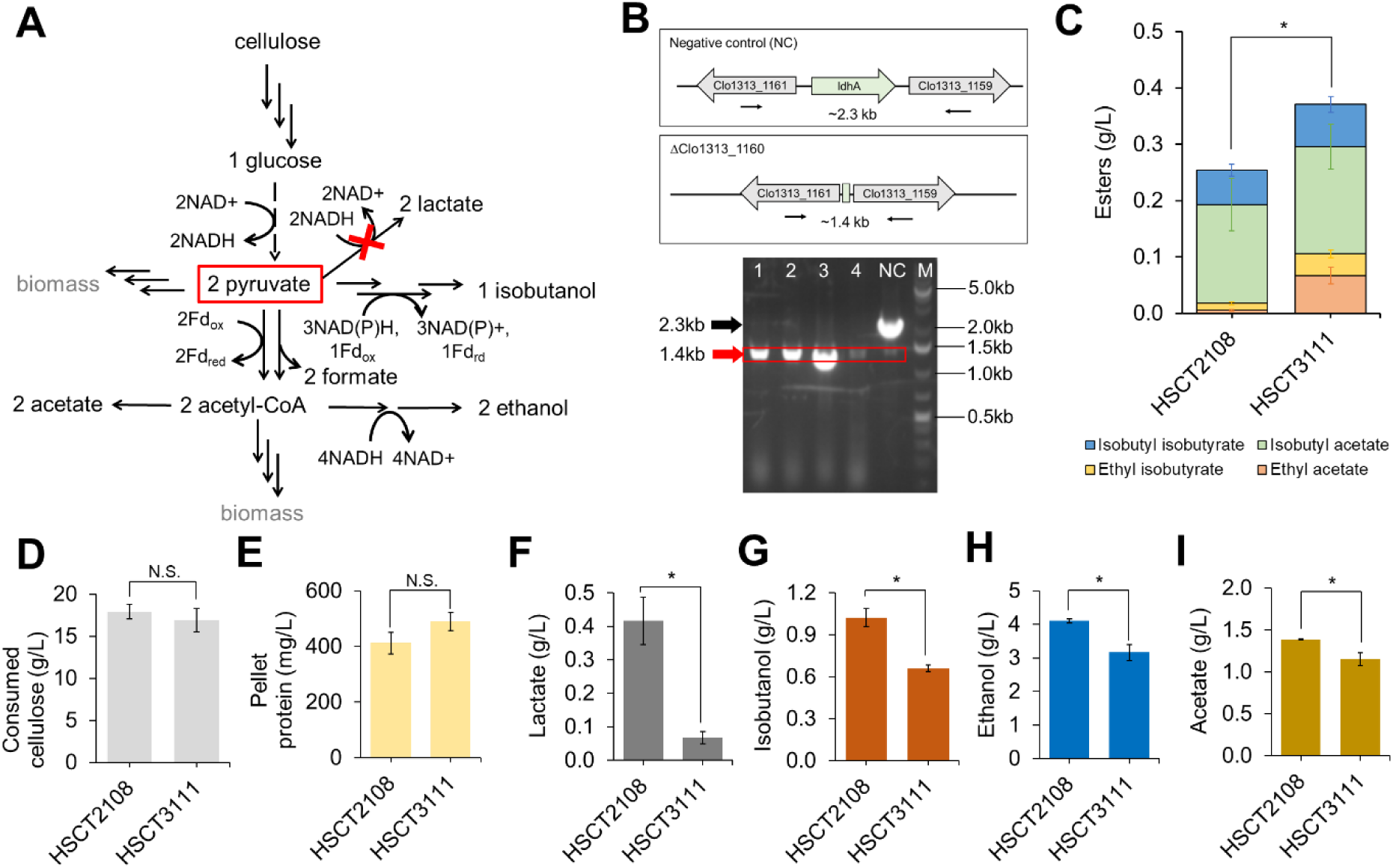
Effect of manipulating the redox-related lactate production pathway and CAT expression on the production and composition of short-chain esters. **(A)** A metabolic map of redox-related central metabolism of *C. thermocellum*. The red symbol “X” indicates deletion of the lactate production pathway. **(B)** Deletion of lactate dehydrogenase (*ldhA, clo1313_1160*) from the *C. thermocellum* chromosome. The red box indicates the expected band size at 1.4 kb upon successful deletion of *ldhA*. The black arrow indicates expected band size at 2.3 kb from a negative control (NC). Lanes 1-4: individual colonies screened by colony PCR, M: DNA ladder. **(C-I)** End point titers of **(C)** short-chain esters, **(D)** consumed cellulose, **(E)** pellet protein, **(F)** lactate, **(G)** isobutanol, **(H)** ethanol, and **(I)** acetate by *C. thermocellum* HSCT2108 and HSCT3111 after 120 h Avicel fermentation. HSCT3111 is derived from HSCT2108 that lacks a *ldhA* gene and expresses CATec3 Y20F under a truncated *gapDH* promoter. Each data point represents a mean ± 1 standard deviation from at least three biological replicates. Abbreviations: N.S., statistically not significant (two tailed Student’s t-test p-value = 0.10); *p-value<0.05 from a two tailed Student’s t-test.

Strain characterization under the Avicel fermentation showed that HSCT3111 had similar cellulose consumption and cell pellet protein like HSCT2108 (Figs. 4D and 4E), indicating that the *ldhA* deletion and CATec3 Y20F overexpression marginally affected cell physiology of HSCT3111. As expected, HSCT3111 produced significantly less lactate than HSCT2108 due to the *ldhA* deletion (Fig. 4F). Minor production of lactate was still observed, indicating there existed a promiscuous lactate dehydrogenase activity in *C. thermocellum,* attributing to the minor lactate dehydrogenase gene encoded by *clo1313_1878*. After 96-hour Avicel fermentation, HSCT3111 (0.38 g/L) completely consumed cellulose and achieved about 1.7-fold increase in the short-chain ester production as compared to HSCT2108 (0.22 g/L) (Fig. 4C) and about 80-fold increase as compared to the initial strain HSCT0102 (0.005 g/L). The enhanced ester production indicated that the truncated *gapDH* promoter effectively overexpressed CATec3 Y20F. We found that HSCT3111 improved ethyl acetate (0.07 g/L) and ethyl isobutyrate (0.04 g/L) production by 4-fold and 2-fold, respectively, while there were no significant differences in isobutyl acetate (0.19 g/L) and isobutyl isobutyrate (0.08 g/L) production between HSCT3111 and HSCT2108. The increase in the short-chain ester production of HSCT3111 was traded off with 34%, 22%, and 17% decrease in isobutanol, ethanol, and acetate production, respectively (Fig. 4G, H, I).

Overall, both deletion of *ldhA* and stable overexpression of CATec3 Y20F improved short-chain ester production. These genetic manipulations marginally affected cell physiology of HSCT3111 in terms of cell growth and substrate consumption as compared to the parent strain HSCT2108.

### Tuning CELF pretreatment conditions to improve production of short-chain esters

To better understand the compatibility of HSCT3111 with CBP using the CELF-pretreated poplar, we investigated the impact of solids produced by different reaction conditions on the short-chain ester production. We produced three different pretreated solids by adjusting the temperature of CELF pretreatment using the steam-jacketed reactor because the scaled-up configuration produced the most biocompatible solids for short-chain ester production compared to the smaller-scale Parr reactor (Fig. 3E). The material was produced from reaction temperatures of 150℃, 160℃, and 170℃ while maintaining the same reaction duration of 15 minutes (Fig. 5A). The severity parameter of CELF pretreatment is a calculated metric based on temperature, time, duration, and acid loading for normalizing and defining the intensity of the pretreatment from different process types. In analyzing the composition of the pretreated solids, all the three conditions produced enriched (>80% w/w) glucan content (Fig. 5B).

**Figure 5.**
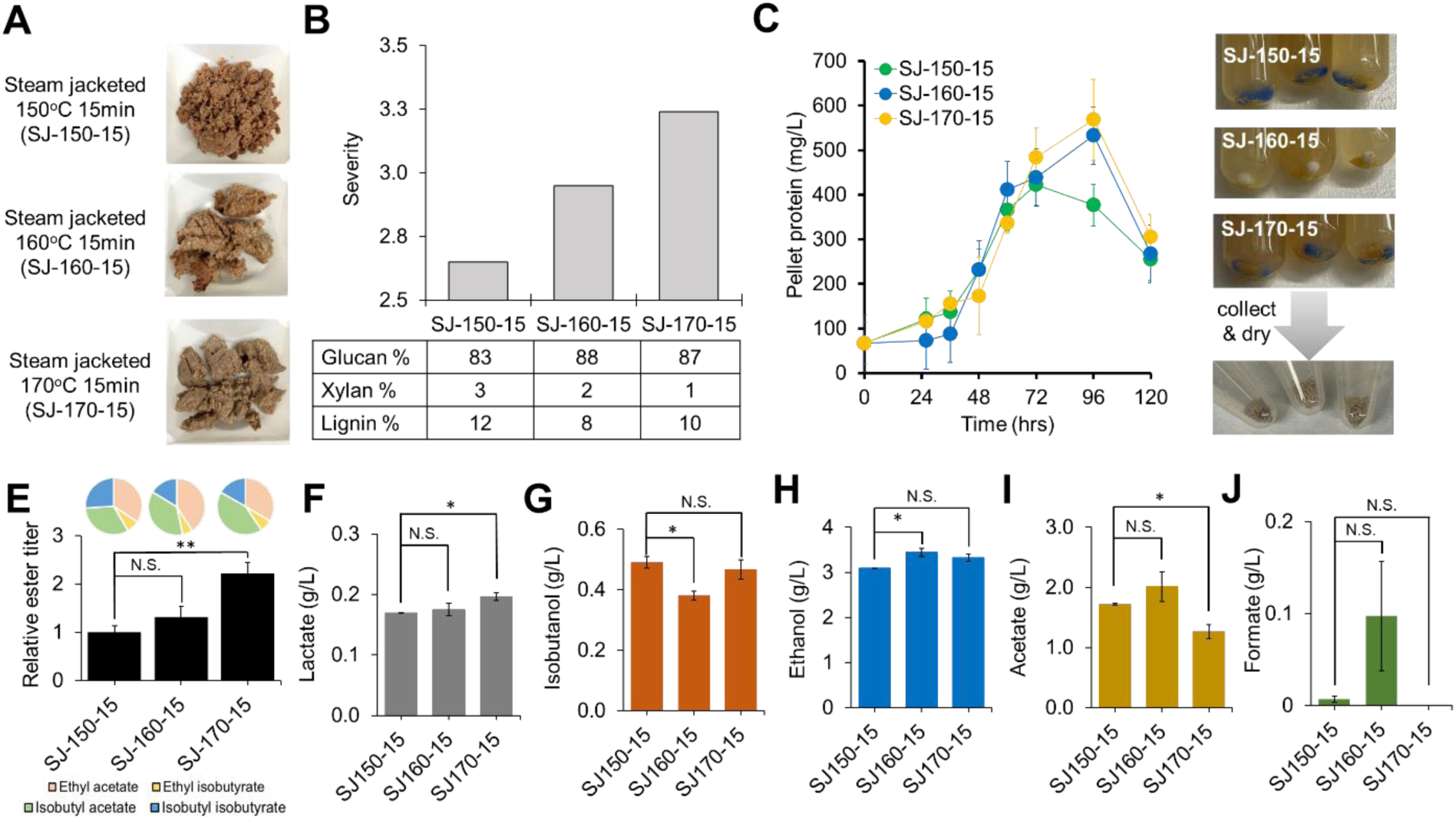
Correlation between the severity of the CELF-pretreated poplar under different process conditions and short-chain ester production by *C. thermocellum* HSCT3111. **(A)** CELF-pretreated poplar under three different conditions (SJ-150-15, SJ-160-15, and SJ-170-15). **(B)** Severity of CELF pretreatment under the different operating conditions. **(C)** Kinetic profiles of pellet protein concentrations (as indicators of cell growth) of *C. thermocellum* HSCT3111 growing on 3 different CELF-pretreated poplar substrates. **(D)** Residual insoluble solids after the CELF-pretreated poplar fermentation. **(E)** Relative ester titers and compositions from three independent fermentation batches of HSCT3111 with at least two biological replicates for each batch after 120-hour fermentation of CELF-pretreated poplar substrates. The pie charts show ester compositions. **(F-J)** End point titers of **(F)** lactate, **(G)** isobutanol, **(H)** ethanol, **(I)** acetate and **(J)** formate by *C. thermocellum* HSCT3111 after 120-hour fermentation of CELF-treated poplar substrates. Each data point represents a mean ± 1 standard deviation from at least three biological replicates. Abbreviations: N.S., statistically not significant (p-value > 0.05); **p-value < 0.01 from a two tailed Student’s t-test; *p-value < 0.05 from two-tailed Students’ t-test.

To examine how the severity of CELF pretreatment affected cell growth and ester production, we next characterized and compared the fermentation of HSCT3111 using the three CELF-pretreated poplar substrates. The results showed that the maximum concentrations of cell pellet proteins were significantly affected by the CELF-pretreated solids (Fig. 5C). Specifically, HSCT3111 yielded the highest cell pellet protein (526 ± 89.7 mg/L) in the SJ-170-15 fermentation using the poplar substrate prepared under the most severe CELF pretreatment (SJ-170-15), followed by the SJ-160-15 fermentation (485 ± 61.9 mg/L) and then the SJ-150-15 fermentation (399.5 ± 33.2 mg/L). At the end of fermentation, HSCT3111 was able to solubilize almost all of the CELF-pretreated solids (Fig. 5D). Because wild-type *C. thermocellum* cannot catabolize xylan and lignin^55^, a mixture of undigested xylan and lignin was expected to be present in the residual pellets. As compared to the SJ-150-15 fermentation (0.04 g/L), the SJ-170-15 fermentation (0.09 g/L) achieved a ∼2.2-fold increase in short-chain ester production (Fig. 5E). From the repetitive fermentation runs, we observed that a short-term growth adaptation of HSCT3111 on the CELF-pretreated solids further enhanced cell growth. Specifically, a longer lag phase of ∼60 hours was observed when the cells were not adapted to the CELF-pretreated poplar substrates at the pre-culture stages (Figs. S4A, S4B). Such delayed fermentation reduced short-chain ester production (Fig. S4C). It should be noted that due to the batch-by-batch performance variations, we performed comparative analysis of the short-chain ester production from three independent batches with at least two biological replicates per batch. Here, the final short-chain ester titers were normalized by the ester titers from the SJ-150-15 fermentation of each batch to compare their production performances (Fig. 5E, S4C).

To examine the effects of CELF severity on fermentative metabolism, we compared the biosynthesis of the major fermentative products including lactate (Fig. 5F), isobutanol (Fig. 5G), ethanol (Fig. 5H), acetate (Fig. 5I), and formate (Fig. 5J). We observed that the production of most of these major fermentative metabolites changed only slightly (i.e., either statistically not significant or less than 10% in differences), suggesting that the cellular redox state and precursor availability might not have been significantly altered by the different CELF-pretreated solids. Acetate production was 26% lower in the SJ-170-15 fermentation than in the SJ-150-15 fermentation (Fig. 5I), likely due to the tradeoff with ester production (Fig. 5E). Similar ethanol and isobutanol production (Figs. 5G, 5H) but different short-chain ester production (Fig. 5E) suggested that the poplar substrates prepared by different CELF pretreatment conditions might have affected the CATec3 Y20F expression level. Since the *gapDH* promoter drove the transcription of CATec3 Y20F, a change in expression of CATec3 Y20F might have been associated with the perturbation of glycolysis to where GapDH belongs. It has been previously reported that lignin and soluble hemicellulose components inhibited *C. thermocellum* cellulosome^56^, reducing solubilization of lignocellulosic biomass^57, 58^. As a result, the CELF-pretreated poplar substrates with different severity (Fig. 5B) might have affected the metabolic capability of *C. thermocellum* to solubilize the solid substrates and the glycolytic fluxes (Figs. 5B, C, D).

## Conclusions

Enabling biological production of biofuels, biochemicals, and biomaterials from lignocellulosic biomass is important for sustainable bioeconomy. A critical challenge is to design a cost-competitive bioprocessing that utilizes renewable and sustainable lignocellulosic biomass feedstocks to produce cost-competitive target products^1^. To tackle the challenge, this study demonstrated, for the first time, the feasibility of a one-step conversion of woody poplar into non-native short-chain esters by engineering *C. thermocellum* to be compatible with CBP coupled with CELF pretreatment. This compatibility required the deletion of CEs, stable overexpression of thermostable and efficient AATs, and rewiring carbon and electron metabolisms of *C. thermocellum* that can utilize the CELF-pretreated poplar substrates to make the desirable short-chain esters.

*C. thermocellum* has the endogenous acetyl-CoA and isobutyryl-CoA pathways that can be rerouted for short-chain ester production. We successfully demonstrated that these endogenous pathways could be activated to produce acetate and isobutyrate esters by deleting CEs and overexpressing thermostable AATs. We found that two CEs (Clo1313_0613 and Clo1313_0693) of *C. thermocellum* not only degraded short-chain esters but also competed for the precursor metabolites due to their promiscuous thioesterase activities. Their deletion was important for short-chain ester biosynthesis while not compromising biomass degradation. Previously, overexpression of the endogenous 2-ketoisovalerate pathway and heterologous ketoacid decarboxylase (*kivD*) in *C. thermocellum* using high cell density, non-growth cultures (i.e., an optical density of 16) was necessary for high isobutanol production ^15^. In contrast, this study demonstrated that it is feasible to activate the endogenous isobutyryl-CoA pathway through the CEs deletion and CATec3 Y20F overexpression to produce isobutanol and associated esters under a typical fermentation condition. We anticipate that future metabolic engineering of *C. thermocellum* to improve ethanol and/or isobutanol production will further increase the short-chain ester synthesis^15, 59, 60^.

From the CBP experiments of *C. thermocellum* using various CELF-pretreated poplar substrates, we found that the short-chain ester production depended on the CELF pretreatment conditions (e.g., reactor configurations and temperatures) that yielded different glucan contents together with minor but inhibitory amounts of xylan and lignin. It has been somewhat controversial whether insoluble lignin interferes with the *C. thermocellum* cell growth and cellulose hydrolysis^55–58, 61, 62^. Our results indicated that the residual xylan and lignin after CELF pretreatment likely inhibited cell growth and ester production (Figs. 3 and 5), and that CBP coupled with higher CELF severity is advantageous to reduce the inhibition from xylan and lignin. Because lignocellulose-derived inhibitors caused substantial changes in proteome and metabolome^63^, a future comprehensive study should analyze the inhibitors from the CELF treated lignocellulose feedstocks. For example, structural changes of glucan or lignin by different CELF severity could also affect the metabolism of *C. thermocellum*^64^. In summary, this study elucidated and enhanced the metabolic compatibility of *C. thermocellum* for conversion of woody poplar into short-chain esters by integrating CBP with CELF pretreatment.

## Materials and Methods

### Strains and plasmids

The list of strains and plasmids of this study are presented in Tables 1 and 2, respectively. Table S1 shows the primers used to construct and confirm the plasmids and strains.

**Table 1:** List of plasmids and strains used in this study.

| Name | Descriptions | Source |
| --- | --- | --- |
| **Plasmids** |  |  |
| pHS0024 | pNW33N derivative, the *cat* gene originated from *Staphylococcus aureus* under the control of *C. thermocellum* gapDH promoter | Ref. 34 |
| pHS0024_F97W | F97W mutant of the CATsa under the control of T7lac promoter, 6X His-tag at N-terminus | Ref. 34 |
| pHS0059_Y20F | A plasmid expressing CATec3 Y20F under the control of *C. thermocellum* gapDH promoter | Ref. 29 |
| pHS005 | pNW33N derivative, The backbone plasmid for gene deletions | Ref. 38 |
| pHS0047 | *clo1313_1160* markerless deletion plasmid | This study |
| pHS0070 | A pHS0059_Y20F derived plasmid expressing CATec3 Y20F under the control of truncated 180bp length *C. thermocellum* gapDH promoter | This study |
| pETDuet-1 | pBR322 ori, Amp<sup>R</sup>, lacI, T7lac promoter | Novagen |
| pET_0613 | Clo1313_0613 cloned at MCS1 site of pETDuet-1. 6X His at N-terminus | Ref. 38 |
| pET_0693 | Clo1313_0693 cloned at MCS1 site of pETDuet-1. Original signal peptides removed and tagged with 6X His at N-terminus | Ref. 38 |
| ***E. coli*** |  |  |
| Top10 | Host for molecular cloning, *mcrA*, $\Delta(mrr-hsdRMS-mcrBC)$, *Phi80lacZ(del)M15*, $\Delta lacX74$, *deoR*, *recA1*, *araD139*, $\Delta(ara-leu)7697$, *galU*, *galK*, *rpsL(SmR)*, *endA1*, *nupG* | Invitrogen |
| BL21 (DE3) | Host for plasmid *in vivo* methylation, *E. coli* B *dcm*, *ompT*, *hsdS(rB-mB-)*, *gal* | Invitrogen |
| BL21-CodonPlus (DE3)- RIPL | Host for the 6XHis-tagged Clo1313_0613 and Clo1313_0693 expression, *E. coli* B F- *ompT hsdS(rB- mB-)* *dcm+* *gal* $\lambda(DE3)$ *endA* *Hte* [*argU proL*] [*argU ileY leuW*], Tet<sup>R</sup>, Cm<sup>R</sup>, Strep/Spec<sup>R</sup> | Agilent |
| ***C. thermocellum*** |  |  |
| M1354 | *C. thermocellum* DSM1313 $\Delta hpt$ | Ref. 14 |
| HSCT2005 | *C. thermocellum* DSM1313 $\Delta hpt$ $\Delta clo1313_0613$ $\Delta clo1313_0693$ | Ref. 68 |
| HSCT0102 | M1354 harboring pHS0024_F97W | Ref. 47 |
| HSCT0103 | M1354 harboring pHS0059_Y20F | This study |
| HSCT2105 | HSCT2005 harboring pHS0024_F97W | Ref. 68 |
| HSCT2108 | HSCT2005 harboring pHS0059_Y20F | Ref. 29 |
| HSCT3009 | *C. thermocellum* DSM1313 $\Delta hpt$ $\Delta clo1313_0613$ $\Delta clo1313_0693$ $\Delta clo1313_1160$ (*ldhA*) | This study |
| HSCT3111 | HSCT2005 harboring pHS0070 | This study |

#### Plasmid construction

Plasmids were constructed by using Gibson DNA assembly and/or ligase-dependent cloning methods. Target DNA fragments were amplified using the Phusion DNA polymerase (cat# F530S, Thermo Fisher Scientific, MA, USA) and purified by DNA purification and gel extraction kits (Omega Biotek, GA, USA). For the ligation-dependent cloning, the backbone plasmids and insert DNA fragments were treated by appropriate restriction enzymes and ligated using a T4 DNA ligase (cat# M0202L, New England Biolabs, MA, USA). Gibson DNA assembly cloning was performed using the purified vector and insert DNA fragments mixed together with the Gibson master mix at 50℃ for one hour^65^. *Escherichia coli* TOP10 was heat-shock transformed after mixed with the assembled DNA and selected on LB agar plates (15 g/L agar) with appropriate antibiotics. All the constructed plasmids were checked by PCR amplification and/or restriction enzyme digestion followed by Sanger sequencing.

#### Strain construction

*C. thermocellum* cells were transformed by electroporation based on the previously described methods^34, 66^. Briefly, after mixing with 1∼2 μg methylated plasmids^67^, cells were electroporated and transferred into a warmed CTFuD liquid medium. For the electroporation, a series of two consecutive exponential pulses were applied using an electroporation system (cat # 45-0651, BTX Technologies Inc., MA, USA). The parameters were set at 1.8 kV, 25 μF, and 350 Ω, which resulted in a pulse duration of 7.0-8.0 milliseconds. After up to 12 hours of recovery, transformed cells were transferred into an anaerobic chamber and plated by mixing cells with a molten CTFuD agar medium inside the chamber. After 3-5 days, colonies were isolated by streaking on a new CTFuD agar plate and used for further studies. For *ldhA* markerless gene deletion, cells after single-crossover event were selected on a CTFuD-NY plate containing thiamphenicol and 5-fluoro-2′-deoxyuradine (FUDR), followed by counterselection for double-crossover event on a CTFuD-NY plate using 8-azahypoxanthine (8AZH)^38, 66^. The cells were consecutively passaged on a new CTFuD plate at least three times to ensure isolation of a single genotypic colony. Deletion of the *ldhA* gene was verified by colony PCR.

### Co-solvent Enhanced Lignocellulosic Fractionation (CELF) pretreatment

#### CELF pretreatment in the Parr Reactor

Downscaled CELF pretreatments were performed in a 1 L Hastelloy Parr® autoclave reactor (236HC Series, Parr Instruments Co., Moline, IL) equipped with a double stacked pitch blade impeller rotated at 200 rpm. The reaction medium contained a mixture of 1:1 water-tetrahydrofuran with 0.05M sulfuric acid. Poplar wood chips were knife-milled through a 1mm particle screen and added to the reactor to achieve a 7.5% (g/g) solid loading in the reactor. The reactions were left to soak overnight at 4°C. The downscaled CELF reactions were ran at 140°C, 150°C and 160°C for a duration of 15, 25, and 35 minutes in order to identify the condition which promoted the highest esters production during CBP. All reactions were maintained at reaction temperature (± 0.5 °C) by submerging the entire reactor and impeller into a 4 kW-heated fluidized sand bath (Model SBL-2D, Techne, Princeton, NJ). The reaction temperature was directly measured by u sing an Inconel-lined submerged K-type thermocouple (Omega Engineering Inc., Stamford, Connecticut). To rapidly arrest the pretreatment, the Parr reactor lifted by a chain-hoist out of the sand bath and immediately submerged into a 50-gal water bath at 25C (room temperature) until the internal temperature dropped to below 35 °C. The reactor was then opened, and the reactant was poured through a ceramic vacuum filter funnel with a glass fiber filter paper (Fisher Scientific, Pittsburgh, PA) to separate the pretreated solids from the pretreated liquor. The vacuum-filtered solids were washed by pouring room temperature DI water through the vacuum filter until the filtrate pH increased to above 6.5. The solids were then collected to be introduced to CBP.

#### CELF pretreatment in Steam-jacketed reactor

Scaled-up CELF pretreatments were performed in an unstirred 1-gal hastelloy steam-assisted reactor. Each reaction was prepared at a total loading of 2800 grams. Poplar wood chips were milled through a 1 mm particle screen and added to the reactor at a 7.5% (g/g) solid loading. A solution containing 1:1 (w/w) water-tetrahydrofuran and 0.05M sulfuric acid was then added and the biomass-solution mixture was allowed to soak overnight. CELF reactions were performed at 150, 160 and 170 °C, all for a15 minute duration. After the reaction, the entire contents of the reactor were released under pressure into a collection tank containing 19 mL of 26 DEG Be ammonium hydroxide solution (∼29%) and partially neutralized to a pH of 3. The collection tank is cooled for 5 minutes under running water. The reaction slurry was then vacuum filtered at room temperature to separate the CELF solids from the liquids. The solids were then further washed with deionized water. The resulting solids were analyzed for compositional differences and were subjected to fermentation utilizing the engineered *C. thermocellum* stain.

### Media and Cell Culturing

*E. coli* strains were grown in lysogeny broth (LB) medium supplemented with antibiotics at concentrations of 30 µg/mL chloramphenicol (Cam) and/or 100 µg/mL ampicillin (Amp) that were used for selection of transformants. *C. thermocellum* strains were cultivated in an anaerobic chamber (Sheldon manufacturing, OR, USA) filled with an anaerobic gas mixture (90% N_2_, 5% CO_2_, 5% H_2_). When culturing *C. thermocellum* strains outside of the chamber, rubber stopper sealed Balch tubes prepared in the chamber were used to maintain anaerobic conditions.

For genetic engineering of *C. thermocellum*, CTFuD or CTFuD-NY medium^66^ supplemented with 10 µg/mL thiamphenicol (Tm) was used. The CTFuD medium contained 2 g/L yeast extract, while the CTFuD-NY medium contained vitamins and trace elements without the yeast extract^66^. Particularly, CTFuD-NY was used instead of CTFuD for counterselection through 8-azahypoxanthine (8-AZH) or 5’-fluoro-2’-deoxyuridine (FuDR). For seed culture, *C. thermocellum* strains were grown in a defined MTC medium with 5 g/L cellobiose as previously described^42^. For ester production, a modified MTC medium (C-MTC) was used as previously described^34^. The C-MTC medium contained (per liter): 2 g urea, 1.5 g ammonium chloride, 10 g 3-(N-morpholino)propanesulfonic acid (MOPS), 2 g sodium citrate tribasic dihydrate, 1.5 g citric acid monohydrate, 1 g sodium sulfate, 0.5 g potassium phosphate monobasic, 1 g cysteine-HCl, 0.2 g calcium chloride, 1 g magnesium chloride hexahydrate, 0.1 g iron (II) chloride tetrahydrate, 2.5 g sodium bicarbonate, 0.02 g pyridoxamine dihydrochloride, 0.004 g p-aminobenzoic acid, 0.002 g biotin, and 0.002 g vitamin B12, and 20 g/L Avicel or other cellulose substrates equivalent to 20 g/L glucan. pH of the C-MTC medium was adjusted to 7.5 and autoclaved for sterilization. Solid LB and CTFuD media additionally contained 15 g/L and 10 g/L of agar, respectively.

#### Lignocellulosic biomass fermentation

Tube-scale cellulose fermentation was performed as previously described^34^. A single colony was inoculated in the MTC medium containing 5 g/L cellobiose and 10 µg/mL Tm and cultivated for 24 hours. For the seed culture adaptation to the CELF-pretreated poplar, the grown cells were passaged to 5 mL MTC medium containing 2.5 g/L cellobiose and 9.5 g/L glucan loading of the CELF-pretreated poplar. The seed culture was periodically mixed by careful pipetting every 4 to 8 hours and incubated for 24 hours at 55 ℃ followed by main culture. For the main culture, 15.2 mL of C-MTC medium was prepared in a Balch tube, and 0.8 mL of overnight cell culture was inoculated in the C-MTC medium in the anaerobic chamber. Then, 4 mL of hexadecane was overlaid on the medium. Each tube contained a small magnetic stirrer bar (10 mm length, 3 mm diameter) to homogenize the insoluble cellulose substrates. The culture was incubated in a water bath connected with a temperature controller set at 55℃ and a magnetic stirrer system. For every 12-24 hours, pH was adjusted with 40-70 μL of 5 M KOH to maintain pH range from 6.4 to 7.5. After the pH adjustment, 800 μL of cell culture and 200 μL of hexadecane layer were sampled for analysis.

#### Analytical methods

##### Cell growth measurement

Cell growth from cellulose fermentation experiments was measured by quantifying pellet protein determined by the Bradford assay as previously described^34^.

##### Residual cellulose quantification from fermentation

Residual cellulose was quantified using phenol-sulfuric acid method with slight modification^34^. Standard curves were determined by using Avicel PH-101 at concentrations of 20 g/L, 10 g/L, 5 g/L, 1 g/L, 0.5 g/L, and 0.1 g/L for every experiment.

##### *In vitro* esterase and thioesterase assay

Clo1313_0613 and Clo1313_0693 were expressed in *E. coli* and His-tag purified as described previously^38^. Esterase activities of the His-tag purified Clo1313_0613 and Clo1313_0693 were measured by quantifying deacetylation rate of p-nitrophenyl acetate (pNPA) on a 96-well plate. The reaction buffer contained 50 mM Tris-HCl at pH 6.8 and 20 mM pNPA in 200 μL of total reaction volume. The reaction started by adding the enzymes at 1 μg/mL concentration, and then an absorbance kinetics at 405 nm was monitored in a microplate reader at 50℃ for 10 minutes. The unit esterase activity was calculated using a standard curve of 0-200 μM p-nitrophenol under the same condition.

Thioesterase activities of the His-tag purified Clo1313_0613 and Clo1313_0693 were measured by DTNB assay using isobutyryl-CoA as a substrate on a 384-well plate. The reaction solution consisted of 50 mM Tris-HCl at pH 6.8, 0.5 mM of isobutyryl-CoA, and 1 mg/mL DTNB in 100% DMSO in 50 μL of total reaction volume. 10 μg/mL of the His-tag purified enzymes were added to the reaction solution and immediately incubated at 50℃ in a plate reader followed by kinetic measurement at 412nm for 30 mins. The reaction rate was calculated using the extinction coefficient from a standard curve of free coenzyme A (MP Biomedicals, OH, USA) under the same condition.

##### Extracellular metabolite analysis

For quantification of extracellular metabolites, an HPLC system (Shimadzu Inc., MD, USA) was used. 800 μL of culture samples were centrifuged at 17,000 xg for 3 minutes and filtered through 0.2 μm pore size syringe filters. The samples were run with 10 mM H_2_SO_4_ at 0.6 mL/min on an Aminex HPX-87H (Biorad Inc., CA, USA) column at 50℃. Concentrations of sugars, organic acids, and alcohols were determined using Refractive index detector (RID) and ultra-violet detector (UVD) at 220 nm.

##### Gas chromatography coupled with mass spectroscopy (GC/MS) analysis

GC (HP 6890, Agilent, CA, USA) equipped with a MS (HP 5973, Agilent, CA, USA) was used to quantify short-chain esters. A Zebron ZB-5 (Phenomenex, CA, USA) capillary column (30 m x 0.25 mm x 0.25 μm) was used with helium as the carrier gas at a flow rate of 0.5 mL/min. The oven temperature program was set as follows: 50℃ initial temperature, 1℃/min ramp up to 58℃, 25℃/min ramp up to 235℃, 50℃/min ramp up to 300℃, and 2-minutes bake-out at 300℃. 1 μL sample was injected into the column with the splitless mode at an injector temperature of 280℃. For the MS system, selected ion mode (SIM) was used to detect and quantify esters with the parameters described previously^34^. As an internal standard, 10 mg/L n-decane were added in initial hexadecane layer and detected with m/z 85, 99, and 113 from 12 to 15-minute retention time range.

#### Compositional analysis of poplar

Compositional analysis of the Poplar before and after CELF pretreatment was done following an established Laboratory Analytical Procedures (LAPs) (version 8-03-2012) from the National Renewable Energy Laboratory (NREL, Golden, CO). After the two-step acid hydrolysis, the resulting solids and liquid were separated using filtering crucibles. The liquid portion was sampled and analyzed against calibration standards and using HPLC Waters Alliance system e2695 (Waters Co., Milford, MA) equipped with an HPX-87H column (Bio-Rad Aminex®, Bio-Rad Laboratories, Hercules, CA) and a Waters Refractive Index Detector 2414 (Waters Co., Milford, MA). The mobile phase run was a 5 mM sulfuric acid set at a flow rate of 0.6 mL/min. Empower® 3 software package (Empower Software Solutions, Newport Beach, CA) was utilized to collect, view, and integrate the resulting chromatographs. The glucan and xylan content were calculated and adjusted against internal sugar standards. To determine the amount of acid insoluble or Klason-lignin, the solid residues after the filtration process were dried and weighed. Finally, ash and other insoluble matter were further quantified by utilizing a muffle furnace ramped to 575 °C to convert the leftover material in the crucibles into ash.

## Acknowledgments

This research was funded by the DOE BER award (DE-SC0022226), the DOE subcontract grant (DE-AC05-000R22725) from the Center of Bioenergy Innovation (the DOE Bioenergy Research Center funded by the Office of Biological and Environmental Research in the DOE Office of Science), and the DOE Joint Genome Institute. The work conducted by the U.S. Department of Energy Joint Genome Institute, a DOE Office of Science User Facility, is supported under Contract No. DE-AC02-05CH11231. The authors would like to thank the Center of Environmental Biotechnology at UTK for using the GC/MS instrument.

## Competing Interests

The authors declare that they have no competing interests.

## Figure Legends

## Supplementary Information

**Supplementary File S1** contains Table S1 and Figures S1, S2, S3, and S4

**Table S1.** A list of primers used in this study.

**Figure S1.** *In vitro* esterase and thioesterase activities for Clo1313_0613 and Clo1313_0693. **(A)** SDS-PAGE picture of His-tag purified Clo1313_0613 (0613) and Clo1313_0693 (0693). L: Protein ladder. The black arrows indicate the expected size of each protein. **(B)** Kinetic profiles of esterase activities of Clo1313_0613 and Clo1313_0693 using pNPA as the substrate. Protein concentration of 0.1 μg/mL was used for the assay. **(C-D)** Kinetic profiles of thioesterase activities of **(C)** Clo1313_0613 and **(D)** Clo1313_0693 using 0.5 mM isobutyryl-CoA as the substrate. Each data point represents mean ± 1 standard deviation from three biological replicates.

**Figure S2.** Kinetic profiles of major metabolites including (A) ethanol, (B) isobutanol, (C) acetate, and (D) lactate from different lignocellulosic biomass fermentation by *C. thermocellum* HSCT2108. Each data point represents a mean ± 1 standard deviation from three biological replicates.

**Figure S3.** Integration of pHS0024 at the chromosomal location of gapDH promoter in *C. thermocellum*. Three biological replicate cultures were PCR screened, indicated as r1, r2, and r3. A *C. thermocellum* containing pNW33N plasmid was used as the negative control (-cont).

**Figure S4.** Fermentation of the CELF-pretreated Poplar with different severity by *C. thermocellum* HSCT3111. **(A-B)** Growth kinetic profiles of HSCT311 **(A)** without seed culture adaptation and **(B)** with 24-hour seed culture adaptation. Each data point represents a mean ± 1 standard deviation from three biological replicates. **(C-E)** End-point titers of **(C)** short-chain esters (n=3), **(D)** ethanol (n=9), and **(E)** isobutanol (n=9). Each data point represents a mean ± 1 standard deviation.

## Notes

### Competing Interest Statement

The authors have declared no competing interest.

## References

(1) Lynd, L. R.; Beckham, G. T.; Guss, A. M.; Jayakody, L. N.; Karp, E. M.; Maranas, C.; McCormick, R. L.; Amador-Noguez, D.; Bomble, Y. J.; Davison, B. H.;, et al. Toward low-cost biological and hybrid biological/catalytic conversion of cellulosic biomass to fuels. Energy & Environmental Science 2022, 15 (3), 938–990, 10.1039/D1EE02540F. DOI: 10.1039/D1EE02540F.

(2) Ragauskas, A. J.; Williams, C. K.; Davison, B. H.; Britovsek, G.; Cairney, J.; Eckert, C. A.; Frederick, W. J., Jr.; Hallett, J. P.; Leak, D. J.; Liotta, C. L.;, et al. The path forward for biofuels and biomaterials. Science (New York, N.Y.) 2006, 311 (5760), 484–489. (acccessed Jan 27).

(3) Himmel, M.; Ding, S.; Johnson, D.; Adney, W.; Nimlos, M.; Brady, J.; Foust, T. Biomass recalcitrance: engineering plants and enzymes for biofuels production. Science 2007, 315 (5813), 804–807.

(4) Lynd, L.; Zyl, W.; McBride, J.; Laser, M. Consolidated bioprocessing of cellulosic biomass: an update. Curr Opin Biotechnol 2005, 16 (5), 577–583.

(5) Olson, D. G.; McBride, J. E.; Shaw, A. J.; Lynd, L. R. Recent progress in consolidated bioprocessing. Curr Opin Biotechnol 2012, 23 (3), 396–405. DOI: 10.1016/j.copbio.2011.11.026.

(6) Lynd, L. R.; Liang, X.; Biddy, M. J.; Allee, A.; Cai, H.; Foust, T.; Himmel, M. E.; Laser, M. S.; Wang, M.; Wyman, C. E. Cellulosic ethanol: status and innovation. Curr Opin Biotechnol 2017, 45, 202–211. DOI: 10.1016/j.copbio.2017.03.008.

(7) Nguyen, T. Y.; Cai, C. M.; Kumar, R.; Wyman, C. E. Co-solvent pretreatment reduces costly enzyme requirements for high sugar and ethanol yields from lignocellulosic biomass. ChemSusChem 2015, 8 (10), 1716–1725.

(8) Bhalla, A.; Cai, C. M.; Xu, F.; Singh, S. K.; Bansal, N.; Phongpreecha, T.; Dutta, T.; Foster, C. E.; Kumar, R.; Simmons, B. A.;, et al. Performance of three delignifying pretreatments on hardwoods: hydrolysis yields, comprehensive mass balances, and lignin properties. Biotechnology for Biofuels 2019, 12 (1), 213. DOI: 10.1186/s13068-019-1546-0.

(9) Patri, A. S.; Mostofian, B.; Pu, Y.; Ciaffone, N.; Soliman, M.; Smith, M. D.; Kumar, R.; Cheng, X.; Wyman, C. E.; Tetard, L.;, et al. A Multifunctional Cosolvent Pair Reveals Molecular Principles of Biomass Deconstruction. Journal of the American Chemical Society 2019, 141 (32), 12545–12557. DOI: 10.1021/jacs.8b10242.

(10) Smith, M. D.; Mostofian, B.; Cheng, X.; Petridis, L.; Cai, C. M.; Wyman, C. E.; Smith, J. C. Cosolvent pretreatment in cellulosic biofuel production: effect of tetrahydrofuran-water on lignin structure and dynamics. Green Chemistry 2016, 18 (5), 1268–1277, 10.1039/C5GC01952D. DOI: 10.1039/C5GC01952D.

(11) Mostofian, B.; Cai, C. M.; Smith, M. D.; Petridis, L.; Cheng, X.; Wyman, C. E.; Smith, J. C. Local Phase Separation of Co-solvents Enhances Pretreatment of Biomass for Bioenergy Applications. Journal of the American Chemical Society 2016, 138 (34), 10869–10878. DOI: 10.1021/jacs.6b03285.

(12) Thomas, V. A.; Donohoe, B. S.; Li, M.; Pu, Y.; Ragauskas, A. J.; Kumar, R.; Nguyen, T. Y.; Cai, C. M.; Wyman, C. E. Adding tetrahydrofuran to dilute acid pretreatment provides new insights into substrate changes that greatly enhance biomass deconstruction by Clostridium thermocellum and fungal enzymes. Biotechnology for Biofuels 2017, 10 (1), 252. DOI: 10.1186/s13068-017-0937-3.

(13) Kothari, N.; Holwerda, E. K.; Cai, C. M.; Kumar, R.; Wyman, C. E. Biomass augmentation through thermochemical pretreatments greatly enhances digestion of switchgrass by Clostridium thermocellum. Biotechnology for Biofuels 2018, 11 (1), 219. DOI: 10.1186/s13068-018-1216-7.

(14) Argyros, D. A.; Tripathi, S. A.; Barrett, T. F.; Rogers, S. R.; Feinberg, L. F.; Olson, D. G.; Foden, J. M.; Miller, B. B.; Lynd, L. R.; Hogsett, D. A.;, et al. High ethanol titers from cellulose by using metabolically engineered thermophilic, anaerobic microbes. Appl Environ Microbiol 2011, 77 (23), 8288–8294. DOI: 10.1128/AEM.00646-11.

(15) Lin, P. P.; Mi, L.; Morioka, A. H.; Yoshino, K. M.; Konishi, S.; Xu, S. C.; Papanek, B. A.; Riley, L. A.; Guss, A. M.; Liao, J. C. Consolidated bioprocessing of cellulose to isobutanol using Clostridium thermocellum. Metab Eng 2015, 31, 44–52. DOI: 10.1016/j.ymben.2015.07.001.

(16) Tian, L.; Conway, P. M.; Cervenka, N. D.; Cui, J.; Maloney, M.; Olson, D. G.; Lynd, L. R. Metabolic engineering of Clostridium thermocellum for n-butanol production from cellulose. Biotechnol Biofuels 2019, 12, 186. DOI: 10.1186/s13068-019-1524-6.

(17) Lee, J. W.; Trinh, C. T. Towards renewable flavors, fragrances, and beyond. Curr Opin Biotechnol 2020, 61, 168–180. DOI: 10.1016/j.copbio.2019.12.017.

(18) Beekwilder, J.; Alvarez-Huerta, M.; Neef, E.; Verstappen, F. W. A.; Bouwmeester, H. J.; Aharoni, A. Functional Characterization of Enzymes Forming Volatile Esters from Strawberry and Banana. Plant Physiology 2004, 135 (4), 1865–1878. DOI: 10.1104/pp.104.042580.

(19) Mason, A. B.; Dufour, J. P. Alcohol acetyltransferases and the significance of ester synthesis in yeast. Yeast 2000, 16 (14), 1287–1298. DOI: 10.1002/1097-0061(200010)16:14<1287::AID-YEA613>3.0.CO;2-I.

(20) Lee, J. W.; Trinh, C. T. Microbial biosynthesis of lactate esters. Biotechnology for Biofuels 2019, 12 (1).

(21) Layton, D. S.; Trinh, C. T. Engineering modular ester fermentative pathways in Escherichia coli. Metabolic Engineering 2014, 26, 77–88. DOI: 10.1016/j.ymben.2014.09.006.

(22) Layton, D. S.; Trinh, C. T. Expanding the modular ester fermentative pathways for combinatorial biosynthesis of esters from volatile organic acids. Biotechnol Bioeng 2016, 113 (8), 1764–1776. DOI: 10.1002/bit.25947.

(23) Rodriguez, G. M.; Tashiro, Y.; Atsumi, S. Expanding ester biosynthesis in *Escherichia coli*. Nature chemical biology 2014, 10 (4), 259–265.

(24) Chacon, M. G.; Kendrick, E. G.; Leak, D. J. Engineering Escherichia coli for the production of butyl octanoate from endogenous octanoyl-CoA. Peerj 2019, 7. DOI: ARTN e6971 10.7717/peerj.6971.

(25) Horton, C. E.; Bennett, G. N. Ester production in E. coli and C. acetobutylicum. Enzyme and microbial technology 2006, 38 (7), 937–943.

(26) Horton, C. E.; Huang, K.-X.; Bennett, G. N.; Rudolph, F. B. Heterologous expression of the Saccharomyces cerevisiae alcohol acetyltransferase genes in Clostridium acetobutylicum and Escherichia coli for the production of isoamyl acetate. Journal of Industrial Microbiology and Biotechnology 2003, 30 (7), 427–432.

(27) Vadali, R.; Horton, C.; Rudolph, F.; Bennett, G.; San, K.-Y. Production of isoamyl acetate in ackA-pta and/or ldh mutants of Escherichia coli with overexpression of yeast ATF2. Applied Microbiology and Biotechnology 2004, 63 (6), 698–704.

(28) Feng, J.; Zhang, J.; Ma, Y.; Feng, Y.; Wang, S.; Guo, N.; Wang, H.; Wang, P.; Jiménez-Bonilla, P.; Gu, Y. Renewable fatty acid ester production in Clostridium. Nature Communications 2021, 12 (1), 1–13.

(29) Seo, H.; Lee, J. W.; Giannone, R. J.; Dunlap, N. J.; Trinh, C. T. Engineering promiscuity of chloramphenicol acetyltransferase for microbial designer ester biosynthesis. Metab Eng 2021, 66, 179–190. DOI: 10.1016/j.ymben.2021.04.005.

(30) Noh, H. J.; Woo, J. E.; Lee, S. Y.; Jang, Y.-S. Metabolic engineering of Clostridium acetobutylicum for the production of butyl butyrate. Applied microbiology and biotechnology 2018, 102 (19), 8319–8327.

(31) Tai, Y.-S.; Xiong, M.; Zhang, K. Engineered biosynthesis of medium-chain esters in Escherichia coli. Metabolic Engineering 2015, 27, 20–28.

(32) Zhu, J.; Lin, J. L.; Palomec, L.; Wheeldon, I. Microbial host selection affects intracellular localization and activity of alcohol-O-acetyltransferase. Microb Cell Fact 2015, 14, 35. DOI: 10.1186/s12934-015-0221-9.

(33) Urit, T.; Li, M.; Bley, T.; Löser, C. Growth of Kluyveromyces marxianus and formation of ethyl acetate depending on temperature. Applied microbiology and biotechnology 2013, 97, 10359–10371.

(34) Seo, H.; Lee, J.-W.; Garcia, S.; Trinh, C. T. Single mutation at a highly conserved region of chloramphenicol acetyltransferase enables isobutyl acetate production directly from cellulose by Clostridium thermocellum at elevated temperatures. Biotechnology for biofuels 2019, 12 (1), 245.

(35) Nancolas, B.; Bull, I. D.; Stenner, R.; Dufour, V.; Curnow, P. Saccharomyces cerevisiae Atf1p is an alcohol acetyltransferase and a thioesterase in vitro. Yeast 2017, 34 (6), 239–251. DOI: 10.1002/yea.3229 From NLM Medline.

(36) Prat, D.; Hayler, J.; Wells, A. A survey of solvent selection guides. Green Chemistry 2014, 16 (10), 4546–4551, 10.1039/C4GC01149J. DOI: 10.1039/C4GC01149J.

(37) Lange, J. P.; Price, R.; Ayoub, P. M.; Louis, J.; Petrus, L.; Clarke, L.; Gosselink, H. Valeric biofuels: a platform of cellulosic transportation fuels. Angew Chem Int Ed Engl 2010, 49 (26), 4479–4483. DOI: 10.1002/anie.201000655.

(38) Seo, H.; Nicely, P. N.; Trinh, C. T. Endogenous carbohydrate esterases of Clostridium thermocellum are identified and disrupted for enhanced isobutyl acetate production from cellulose. Biotechnol Bioeng 2020, 117 (7), 2223–2236. DOI: 10.1002/bit.27360.

(39) Rydzak, T.; Levin, D. B.; Cicek, N.; Sparling, R. End-product induced metabolic shifts in Clostridium thermocellum ATCC 27405. Appl Microbiol Biotechnol 2011, 92 (1), 199–209. DOI: 10.1007/s00253-011-3511-0 From NLM Medline.

(40) Rydzak, T.; Grigoryan, M.; Cunningham, Z. J.; Krokhin, O. V.; Ezzati, P.; Cicek, N.; Levin, D. B.; Wilkins, J. A.; Sparling, R. Insights into electron flux through manipulation of fermentation conditions and assessment of protein expression profiles in Clostridium thermocellum. Applied Microbiology and Biotechnology 2014, 98 (14), 6497–6510. DOI: 10.1007/s00253-014-5798-0.

(41) Sander, K.; Wilson, C. M.; Rodriguez, M.; Klingeman, D. M.; Rydzak, T.; Davison, B. H.; Brown, S. D. Clostridium thermocellum DSM 1313 transcriptional responses to redox perturbation. Biotechnology for Biofuels 2015, 8 (1), 211. DOI: 10.1186/s13068-015-0394-9.

(42) Thompson, R. A.; Layton, D. S.; Guss, A. M.; Olson, D. G.; Lynd, L. R.; Trinh, C. T. Elucidating central metabolic redox obstacles hindering ethanol production in Clostridium thermocellum. Metab Eng 2015, 32, 207–219. DOI: 10.1016/j.ymben.2015.10.004.

(43) Thompson, R. A.; Trinh, C. T. Overflow metabolism and growth cessation in Clostridium thermocellum DSM1313 during high cellulose loading fermentations. Biotechnol Bioeng 2017, 114 (11), 2592–2604. DOI: 10.1002/bit.26374.

(44) Sista Kameshwar, A. K.; Qin, W. Understanding the structural and functional properties of carbohydrate esterases with a special focus on hemicellulose deacetylating acetyl xylan esterases. Mycology 2018, 9 (4), 273–295. DOI: 10.1080/21501203.2018.1492979.

(45) Lo, Y. C.; Lin, S. C.; Shaw, J. F.; Liaw, Y. C. Substrate specificities of Escherichia coli thioesterase I/protease I/lysophospholipase L1 are governed by its switch loop movement. Biochemistry 2005, 44 (6), 1971–1979. DOI: 10.1021/bi048109x.

(46) Lescic Asler, I.; Ivic, N.; Kovacic, F.; Schell, S.; Knorr, J.; Krauss, U.; Wilhelm, S.; Kojic-Prodic, B.; Jaeger, K. E. Probing enzyme promiscuity of SGNH hydrolases. Chembiochem 2010, 11 (15), 2158–2167. DOI: 10.1002/cbic.201000398.

(47) Seo, H.; Lee, J. W.; Garcia, S.; Trinh, C. T. Single mutation at a highly conserved region of chloramphenicol acetyltransferase enables isobutyl acetate production directly from cellulose by Clostridium thermocellum at elevated temperatures. Biotechnol Biofuels 2019, 12, 245. DOI: 10.1186/s13068-019-1583-8.

(48) Patri, A. S.; Mostofian, B.; Pu, Y.; Ciaffone, N.; Soliman, M.; Smith, M. D.; Kumar, R.; Cheng, X.; Wyman, C. E.; Tetard, L.;, et al. A Multifunctional Cosolvent Pair Reveals Molecular Principles of Biomass Deconstruction. J Am Chem Soc 2019, 141 (32), 12545–12557. DOI: 10.1021/jacs.8b10242 From NLM Medline.

(49) Cai, C. M.; Zhang, T.; Kumar, R.; Wyman, C. E. THF co-solvent enhances hydrocarbon fuel precursor yields from lignocellulosic biomass. GREEN CHEMISTRY 2013, 15 (11), 3140–3145, 10.1039/C3GC41214H. DOI: 10.1039/c3gc41214h.

(50) Nguyen, T. Y.; Cai, C. M.; Osman, O.; Kumar, R.; Wyman, C. E. CELF pretreatment of corn stover boosts ethanol titers and yields from high solids SSF with low enzyme loadings. Green Chemistry 2016, 18 (6), 1581–1589, 10.1039/C5GC01977J. DOI: 10.1039/C5GC01977J.

(51) Olson, D. G.; Maloney, M.; Lanahan, A. A.; Hon, S.; Hauser, L. J.; Lynd, L. R. Identifying promoters for gene expression in Clostridium thermocellum. Metabolic Engineering Communications 2015, 2, 23–29. DOI: https://doi.org/10.1016/j.meteno.2015.03.002.

(52) Tian, L.; Perot, S. J.; Hon, S.; Zhou, J.; Liang, X.; Bouvier, J. T.; Guss, A. M.; Olson, D. G.; Lynd, L. R. Enhanced ethanol formation by Clostridium thermocellum via pyruvate decarboxylase. Microb Cell Fact 2017, 16 (1), 171. DOI: 10.1186/s12934-017-0783-9.

(53) Kim, S. K.; Groom, J.; Chung, D.; Elkins, J.; Westpheling, J. Expression of a heat-stable NADPH-dependent alcohol dehydrogenase from Thermoanaerobacter pseudethanolicus 39E in Clostridium thermocellum 1313 results in increased hydroxymethylfurfural resistance. Biotechnol Biofuels 2017, 10, 66. DOI: 10.1186/s13068-017-0750-z.

(54) Fujitani, Y.; Yamamoto, K.; Kobayashi, I. Dependence of frequency of homologous recombination on the homology length. Genetics 1995, 140 (2), 797–809.

(55) Demain, A. L.; Newcomb, M.; Wu, J. H. Cellulase, clostridia, and ethanol. Microbiol Mol Biol Rev 2005, 69 (1), 124–154. DOI: 10.1128/MMBR.69.1.124-154.2005 From NLM Medline.

(56) Chen, C.; Qi, K.; Chi, F.; Song, X.; Feng, Y.; Cui, Q.; Liu, Y. J. Dissolved xylan inhibits cellulosome-based saccharification by binding to the key cellulosomal component of Clostridium thermocellum. Int J Biol Macromol 2022, 207, 784–790. DOI: 10.1016/j.ijbiomac.2022.03.158 From NLM Medline.

(57) Beri, D.; Herring, C. D.; Blahova, S.; Poudel, S.; Giannone, R. J.; Hettich, R. L.; Lynd, L. R. Coculture with hemicellulose-fermenting microbes reverses inhibition of corn fiber solubilization by Clostridium thermocellum at elevated solids loadings. Biotechnol Biofuels 2021, 14 (1), 24. DOI: 10.1186/s13068-020-01867-w From NLM PubMed-not-MEDLINE.

(58) Kubis, M. R.; Holwerda, E. K.; Lynd, L. R. Declining carbohydrate solubilization with increasing solids loading during fermentation of cellulosic feedstocks by Clostridium thermocellum: documentation and diagnostic tests. Biotechnol Biofuels Bioprod 2022, 15 (1), 12. DOI: 10.1186/s13068-022-02110-4 From NLM PubMed-not-MEDLINE.

(59) Holwerda, E. K.; Olson, D. G.; Ruppertsberger, N. M.; Stevenson, D. M.; Murphy, S. J. L.; Maloney, M. I.; Lanahan, A. A.; Amador-Noguez, D.; Lynd, L. R. Metabolic and evolutionary responses of Clostridium thermocellum to genetic interventions aimed at improving ethanol production. Biotechnol Biofuels 2020, 13, 40. DOI: 10.1186/s13068-020-01680-5.

(60) Hon, S.; Holwerda, E. K.; Worthen, R. S.; Maloney, M. I.; Tian, L.; Cui, J.; Lin, P. P.; Lynd, L. R.; Olson, D. G. Expressing the Thermoanaerobacterium saccharolyticum pforA in engineered Clostridium thermocellum improves ethanol production. Biotechnol Biofuels 2018, 11, 242. DOI: 10.1186/s13068-018-1245-2 From NLM PubMed-not-MEDLINE.

(61) Lynd, L. R.; Grethlein, H. E.; Wolkin, R. H. Fermentation of Cellulosic Substrates in Batch and Continuous Culture by Clostridium thermocellum. Appl Environ Microbiol 1989, 55 (12), 3131–3139. DOI: 10.1128/aem.55.12.3131-3139.1989 From NLM PubMed-not-MEDLINE.

(62) Avgerinos, G. C.; Wang, D. I. Selective solvent delignification for fermentation enhancement. Biotechnol Bioeng 1983, 25 (1), 67–83. DOI: 10.1002/bit.260250107 From NLM PubMed-not-MEDLINE.

(63) Poudel, S.; Giannone, R. J.; Rodriguez, M., Jr.; Raman, B.; Martin, M. Z.; Engle, N. L.; Mielenz, J. R.; Nookaew, I.; Brown, S. D.; Tschaplinski, T. J.;, et al. Integrated omics analyses reveal the details of metabolic adaptation of Clostridium thermocellum to lignocellulose-derived growth inhibitors released during the deconstruction of switchgrass. Biotechnol Biofuels 2017, 10, 14. DOI: 10.1186/s13068-016-0697-5 From NLM PubMed-not-MEDLINE.

(64) Wang, Y.-Y.; Sengupta, P.; Scheidemantle, B.; Pu, Y.; Wyman, C. E.; Cai, C. M.; Ragauskas, A. J. Effects of CELF Pretreatment Severity on Lignin Structure and the Lignin-Based Polyurethane Properties. Frontiers in Energy Research 2020, 8, Original Research. DOI: 10.3389/fenrg.2020.00149.

(65) Gibson, D. G.; Young, L.; Chuang, R. Y.; Venter, J. C.; Hutchison, C. A., 3rd; Smith, H. O. Enzymatic assembly of DNA molecules up to several hundred kilobases. Nat Methods 2009, 6 (5), 343–345. DOI: 10.1038/nmeth.1318.

(66) Olson, D. G.; Lynd, L. R. Transformation of Clostridium thermocellum by electroporation. Methods Enzymol 2012, 510, 317–330. DOI: 10.1016/B978-0-12-415931-0.00017-3.

(67) Guss, A. M.; Olson, D. G.; Caiazza, N. C.; Lynd, L. R. Dcm methylation is detrimental to plasmid transformation in Clostridium thermocellum. Biotechnol Biofuels 2012, 5 (1), 30. DOI: 10.1186/1754-6834-5-30 From NLM PubMed-not-MEDLINE.

(68) Seo, H.; Nicely, P. N.; Trinh, C. T. Endogenous carbohydrate esterases of Clostridium thermocellum are identified and disrupted for enhanced isobutyl acetate production from cellulose. Biotechnol Bioeng 2020. DOI: 10.1002/bit.27360.

