## Supplementary Information for "Consolidated Bioprocessing of Lignocellulosic Biomass Poplar to Produce Short-Chain Esters by *Clostridium thermocellum*"

**Table S1.** A list of primers used in this study. The underlined letters indicate restriction enzyme recognition sites.

| Primer name | Primer sequence (5' to 3') | Description |
|---|---|---|
| HS452 | CTCTGAGCTCCCGGAAGATATGCCTGC | *clo1313_1160* deletion upstream forward |
| HS453 | GAAAAGGAGAACTTATTCCTTCCTCCTTTATCAATAAC | *clo1313_1160* deletion upstream reverse |
| HS454 | ATAAAGGAGGAAGGAATAAGTTCTCCTTTTCTTTTATGA | *clo1313_1160* deletion downstream forward |
| HS455 | CTCTGGATCCATCTTGACAGTATTTCGGATTC | *clo1313_1160* deletion downstream reverse |
| HS456 | TTGTACACGGCCGCATAATCGCCAACCAAAAAGAAGGCG | *clo1313_1160* deletion intermediate reverse |
| HS457 | GCAGCCTAGGTTAATTAAGCTGCGCTCATATATCTAGTGTTTTTATTATTTC | *clo1313_1160* deletion intermediate reverse |
| HS83 | AGCATCGGCATCATCTC | *clo1313_1160* deletion check primer |
| HS86 | CAAGGGGTTTTCGTCGTG | *clo1313_1160* deletion check primer |
| DuetDown-1 | GATTATGCGGCCGTGTACAA | MCS2 region backbone amplification primer to assemble with *clo1313_1160* intermediate fragment |
| HS2 | GCGCAGCTTAATTAACCTAGGCTGC | MCS2 region backbone amplification primer to assemble with *clo1313_1160* intermediate fragment |
| HS606 | GGATGGATTTTAAAAAGCTGGATACT | Primer to check chromosomal integration of pHS0024 |
| HS607 | TAACGCGCTAAACGTCTCTGTTTCC | Primer to check chromosomal integration of pHS0024 |
| HS793 | ATCATGGTCATAGATTTTTTATCTAAACTATTGAAAATGAAA | PgapDH truncation primer |
| HS794 | CAATAGTTTAGATAAAAAATCTATGACCATGATTACGCC | PgapDH truncation primer |

**Figure S1.** *In vitro* esterase and thioesterase activities for Clo1313_0613 and Clo1313_0693. **(A)** SDS-PAGE picture of His-tag purified Clo1313_0613 (0613) and Clo1313_0693 (0693). L: Protein ladder. The black arrows indicate the expected size of each protein. **(B)** Kinetic profiles of esterase activities of Clo1313_0613 and Clo1313_0693 using pNPA as the substrate. Protein concentration of 0.1 µg/mL was used for the assay. **(C-D)** Kinetic profiles of thioesterase activities of **(C)** Clo1313_0613 and **(D)** Clo1313_0693 using 0.5 mM isobutyryl-CoA as the substrate. Each data point represents mean ± 1 standard deviation from three biological replicates.

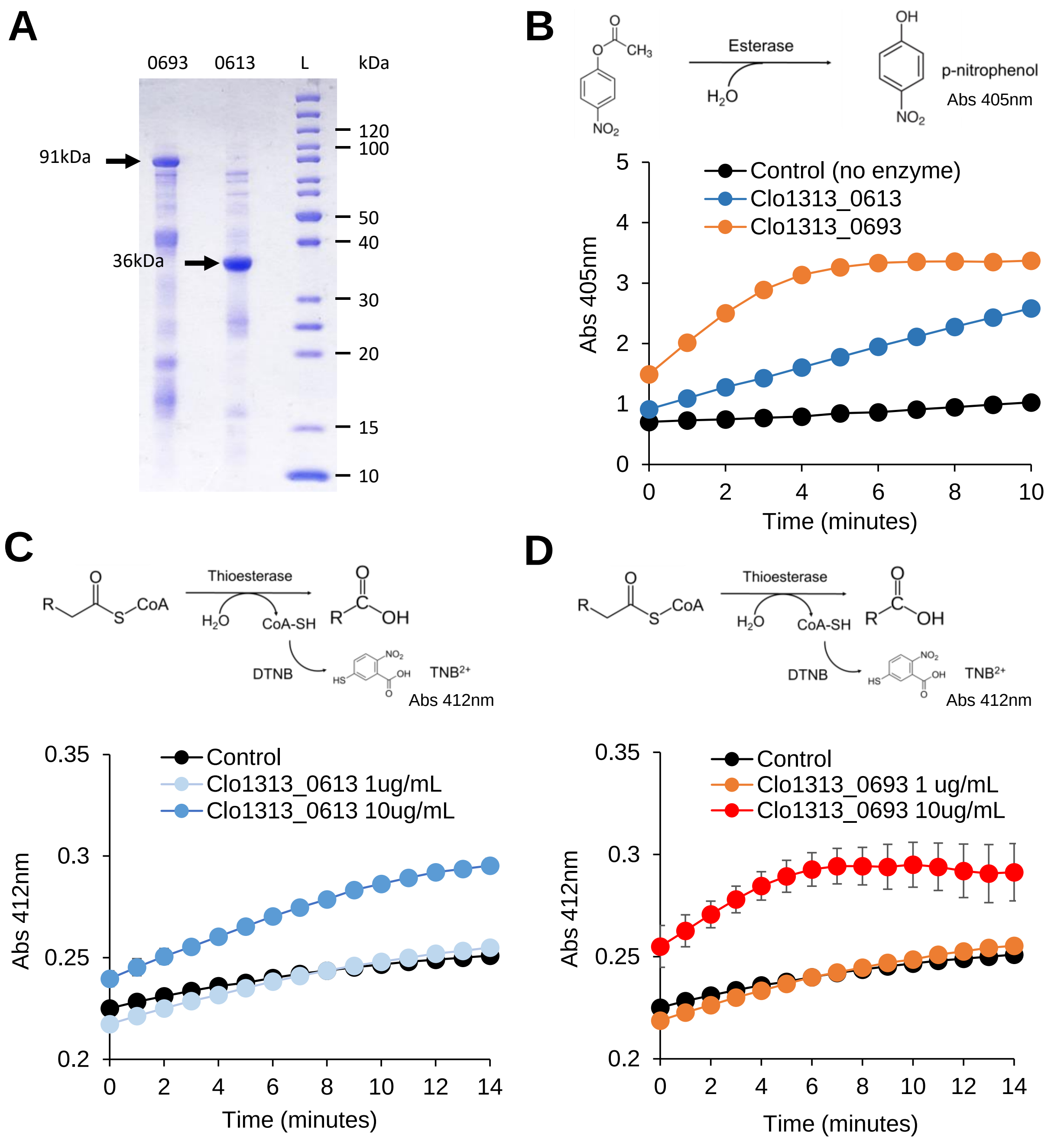

**Figure S2.** Kinetic profiles of major metabolites including **(A)** ethanol, **(B)** isobutanol, **(C)** acetate, and **(D)** lactate from different lignocellulosic biomass fermentation by *C. thermocellum* HSCT2108. Each data point represents a mean ± 1 standard deviation from three biological replicates.

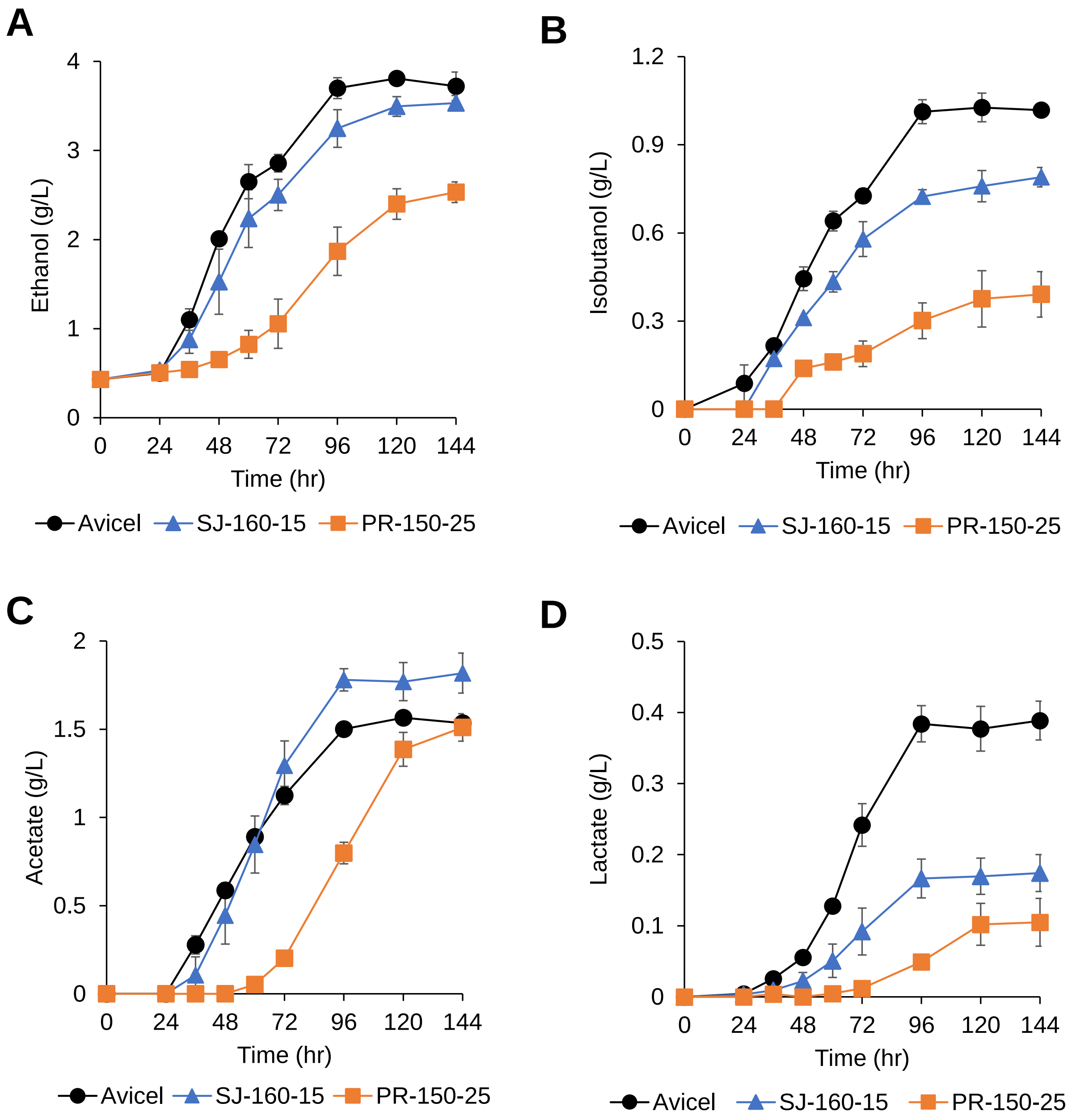

**Figure S3.** Integration of pHS0024 at the chromosomal location of gapDH promoter in *C. thermocellum*. Three biological replicate cultures were PCR screened, indicated as r1, r2, and r3. A *C. thermocellum* containing pNW33N plasmid was used as the negative control (-cont).

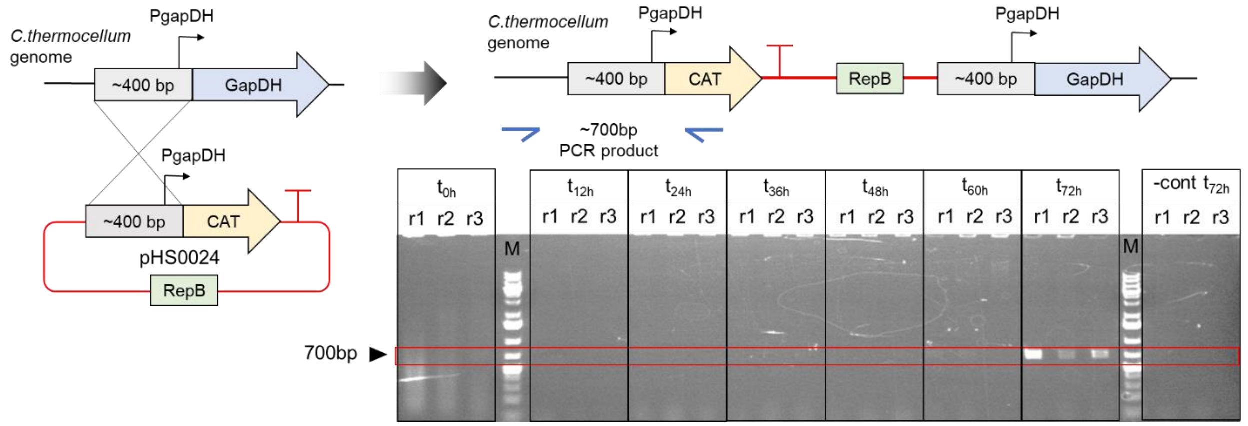

**Figure S4.** Fermentation of the CELF-pretreated Poplar with different severity by *C. thermocellum* HSCT3111. **(A-B)** Growth kinetic profiles of HSCT311 **(A)** without seed culture adaptation and **(B)** with 24-hour seed culture adaptation. Each data point represents a mean ± 1 standard deviation from three biological replicates. **(C-E)** End-point titers of **(C)** short-chain esters (n=3), **(D)** ethanol (n=9), and **(E)** isobutanol (n=9). Each data point represents a mean ± 1 standard deviation.

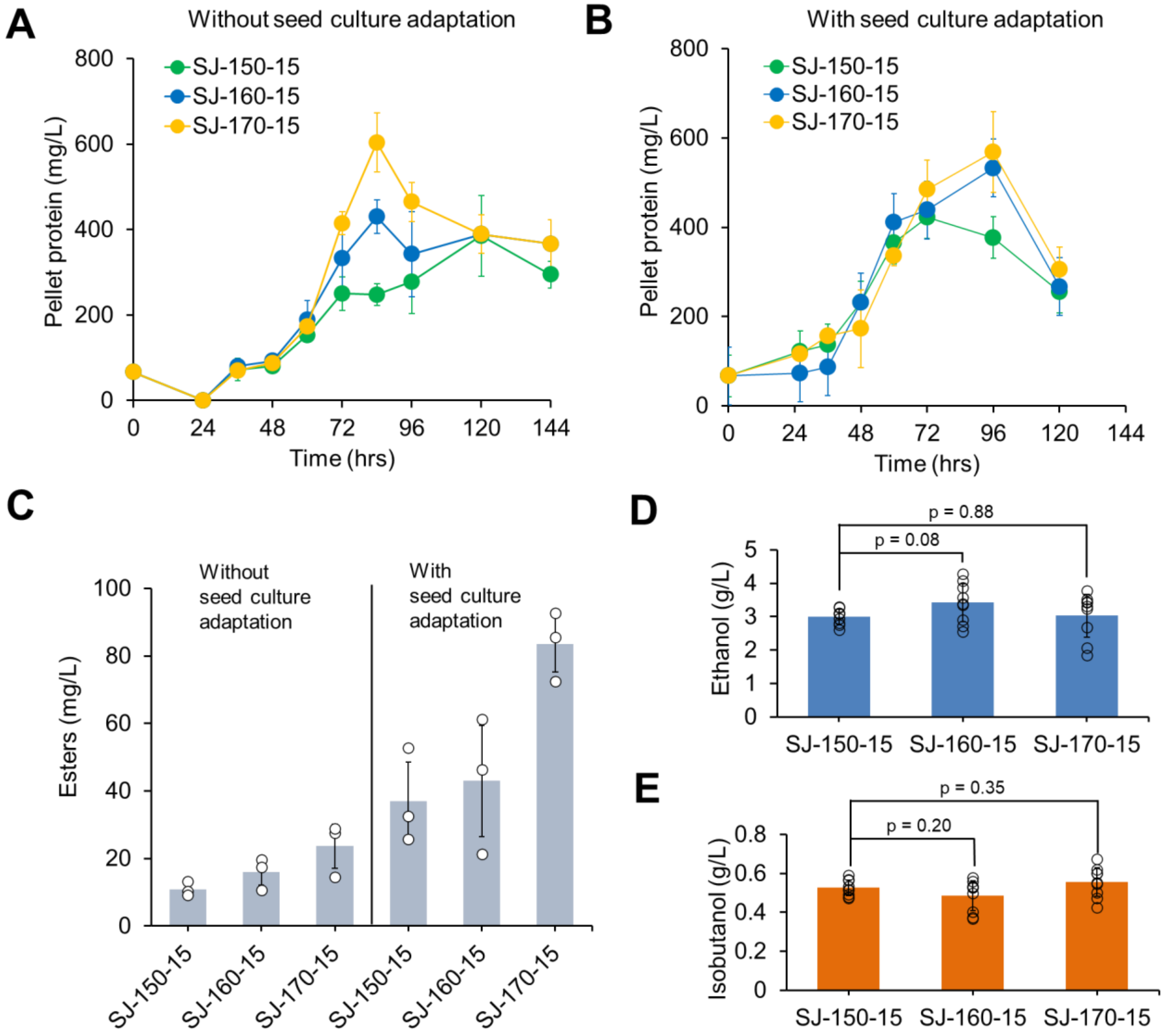
